# MtrAB activates ectoine production and triggers sporulation in response to osmotic stress in *Streptomyces venezuelae*

**DOI:** 10.64898/2025.12.19.695424

**Authors:** Ainsley D.M. Beaton, Rebecca Devine, Neil A. Holmes, Nicolle F. Som, Lucas Balis, Katie Noble, Rhea Stringer, Martin Rejzek, Gerhard Saalbach, Barrie Wilkinson, Matthew I. Hutchings

## Abstract

The MtrAB two-component system is a master regulator of antibiotic biosynthesis in *Streptomyces* species. MtrA is also required for sporulation under certain growth conditions, which means that on some growth media Δ*mtrA* mutant colonies do not produce aerial hyphae or spores. These mutants are referred to as (conditionally) bald because they lack the hairy appearance of wild-type colonies. Here we report that *S. venezuelae* NRRL B-65442 Δ*mtrA* is bald on R2YE agar but sporulates normally on MYM agar and we demonstrate that this is caused by the presence of 10.2% sucrose in R2YE. Consistent with this, we found that adding 10.2% sucrose to MYM agar also inhibits sporulation of the Δ*mtrA* mutant. Proteomics combined with DNA binding studies revealed that MtrA directly activates the expression of the key developmental regulator genes *bldM* and *whiI* on R2YE but not MYM agar. BldM and WhiI work together to activate genes required for aerial hyphae production and sporulation. Crucially, over-expression of *bldM-whiI* in the Δ*mtrA* mutant restored normal sporulation in the presence of 10.2% sucrose. We hypothesised that MtrAB must sense and respond to osmotic stress and consistent with this we found that MtrA directly activates biosynthesis of the compatible solute and osmoprotectant ectoine on growth media containing 10.2% sucrose. We propose a model in which high concentrations of sucrose induce osmotic stress in *Streptomyces* species and this activates MtrAB. The response regulator MtrA then directly activates expression of the *ectABCD* operon to switch on ectoine biosynthesis and expression of *bldM* and *whiI* to trigger entry into sporulation.

**Impact statement:** *Streptomyces* species have complex developmental life cycles that start with spore germination and outgrowth of an actively growing substrate mycelium. Environmental stresses including nutrient starvation and osmotic stress trigger the production of aerial hyphae that undergo cell division to form spores. These spores are more resistant to environmental stresses and can stay viable in the soil until conditions improve. Here we show that MtrAB is responsible for sensing and responding to osmotic stress and we demonstrate that it directly controls the biosynthesis of ectoine and the developmental transition to sporulation through direct activation of BldM and WhiI. This work provides new insight into how environmental cues control the development of *Streptomyces* bacteria.

**Data summary:** The *S. venezuelae* NRRL B-65442 genome sequence is available at NCBI (Reference Sequence NZ_CP018074.1) and can be viewed at http://strepDB.streptomyces.org.uk (select vnz chromosome from the drop-down list). The ChIP-seq data (GEO accession number CP018074) used to map MtrA binding sites on the *S. venezuelae* NRRL B-65442 chromosome and the differential RNA-seq data (accession number GSE81104) used to map global transcript start sites throughout the life cycle were generated for earlier studies (1,2). The tandem mass tag proteomics data are available via ProteomeXchange with identifier PXD069151. The ReDCaT SPR data is included in the manuscript and the supplementary information. Protocols are freely available at http://actinobase.org (3) and strains and plasmids are available from http://streptomyces.org.uk/strepstrains. **The authors confirm all supporting data, code and protocols have been provided within the article or through supplementary data files.**

## Introduction

The genus *Streptomyces* comprises >1100 verified species whose specialised metabolites form the basis of around 55% of clinically used antibiotics (4,5). They are ubiquitous in soils where they play an important role in the turnover of organic material and are enriched in the rhizosphere and endosphere of many different plants (6–9). Their complex developmental life cycles include hyphal growth and sporulation, and these are often linked to the production of bioactive specialised metabolites (4,10). These structurally diverse specialised metabolites play important roles in their interactions with other soil organisms, including microbes, insects, and plants (11,12). Given the diversity of environments they colonise, it is perhaps not surprising that more than 20% of the genes in their large, linear (7-12Mbp) genomes encode transcription factors and signal transduction systems, while up to 10% of their genomes are dedicated to specialised metabolism (13).

This study focuses on the model organism *Streptomyces venezuelae* NRRL B-65442 (5), which encodes 57 two-component signal transduction systems (TCS), with 15 of these conserved throughout the genus (14). TCSs allow bacteria to sense and respond to extracellular signals, and the MtrAB system is highly conserved both within *Streptomyces* species and across the wider phylum Actinomycetota (formerly Actinobacteria) (15). The *mtrAB* operon also includes the *lpqB* gene, encoding a lipoprotein which has been shown to modulate MtrB sensor kinase activity in *Mycobacterium smegmatis* (15,16). The response regulator MtrA was first identified in mycobacteria (<u>M</u>ycobacterial transcriptional regulator <u>A</u>) and has been reported to be essential in *M. tuberculosis* (17) where it coordinates DNA replication with cell division (18–20). In *Corynebacterium glutamicum* MtrAB senses osmotic stress and controls a regulon of genes encoding products involved in the osmotic stress response, including the proline transporter ProP (21). In *Streptomyces* species, MtrA has been implicated in the regulation of antibiotic production and development, including sporulation (22), and binds to at least some of the same sites as the atypical response regulator GlnR to coordinate the regulation of nitrogen metabolism (23,24). MtrA has also been implicated in the regulation of antibiotic production in rare actinomycetes, including erythromycin in *Saccharopolyspora erythraea* (25,26). MtrA directly regulates the production of antibiotics in *S. coelicolor* and *S. venezuelae* and coordinates the production of chloramphenicol with sporulation in *S. venezuelae* NRRL B-65442, such that loss of MtrA activity leads to constitutive high-level production of this antibiotic (1,27,28). MtrA also represses the production of avermectin in *Streptomyces avermitilis* while activating morphological differentiation (29). Thus, modulation of MtrAB activity can be used to increase the production of antibiotics in *Streptomyces* species and other closely related filamentous actinomycetes. Consistent with this, MtrA binding sites have been identified within biosynthetic gene clusters (BGCs) for medically important compounds, including gentamicin, rifamycin, oxytetracycline, daptomycin and streptomycin (28). It appears that MtrA directly controls antibiotic production in these bacteria, but we do not understand how MtrA controls development or why Δ*mtrA* mutants are conditionally bald. This means they form aerial hyphae and spores on some growth media but not others and the reasons for this are not known.

In this work we set out to address this question by comparing MtrA-dependent developmental gene expression in the wild-type and Δ*mtrA* mutant strains grown on R2YE agar (without glucose), where Δ*mtrA* is bald, and MYM agar where both strains sporulate normally. R2YE agar contains 10.2% sucrose, and we demonstrate that omitting sucrose from R2YE enables the Δ*mtrA* mutant to sporulate while adding 10.2% sucrose to MYM agar prevents sporulation. Thus, high sucrose concentrations appear to activate MtrAB activity, most likely by inducing osmotic stress, and this triggers sporulation such that an Δ*mtrA* mutant cannot sporulate under these conditions. Next, we used surface plasmon resonance (SPR) to identify the exact MtrA binding sites at the promoters of the developmental genes *adpA*, *bldM*, *dnaA*, *filP*, *ssgB* and *whiI* which were previously identified as MtrA targets using ChIP-seq (1) (Figure S1). Quantitative tandem mass tag (TMT) proteomics showed that levels of the atypical response regulators BldM and WhiI are reduced 4-fold and 7-fold, respectively, in the Δ*mtrA* mutant grown on R2YE agar but not on MYM agar while the other gene products were unaffected. Homodimeric BldM controls the expression of genes required for the development of aerial hyphae while heterodimeric BldM-WhiI controls the expression of genes required for cell division of the aerial hyphae into spores (30). Over-expression of *bldM-whiI* restored sporulation to the Δ*mtrA* mutant under osmotic stress conditions. We thus hypothesise that MtrAB senses and responds to osmotic stress by triggering entry into sporulation. Consistent with this we show that MtrA also directly activates expression of the *ectABCD* operon which encodes the biosynthetic pathway for the osmoprotectant ectoine (31). We further show that ectoine is produced at elevated levels on MYM and R2YE agar containing 10.2% sucrose and that this is dependent on MtrA. Our data support a model in which high sucrose concentrations trigger an osmotic stress response that is mediated by MtrAB and results in the production of ectoine and spores, thus ensuring survival of these bacteria until conditions improve.

## Methods

For detailed written and video protocols visit http://actinobase.org (3).

### Strains, plasmids and primers

All the bacterial strains used in this work are listed in Table 1 and plasmids are listed in Table 2. Those that were generated for this work were constructed as follows: DNA fragments containing 25–35 nucleotide overlapping regions were assembled into digested DNA vectors using the exonuclease-based Gibson Assembly (NEB). A standard 3:1 ratio of insert to vector was used and assembly performed at 50 °C for 1 hour in a thermocycler. All the primers used for cloning in this work are listed in Table S1.

**Table 1.** Bacterial strains used in this work.

| Bacterial strain | Description | Supplier and/or reference |
| --- | --- | --- |
| *E. coli* DH5alpha | *E. coli* strain used for cloning | ThermoFisher |
| *E. coli* Top10 | F<sup>−</sup> *mcrA* $\Delta$(*mrr-hsdRMS-mcrBC*)<br>Φ80*lacZ*ΔM15 Δ*lacX74* *recA1* *araD139*<br>$\Delta$(*ara leu*) 7697 *galU* *galK* *rpsL* (StrR)<br>*endA1* *nupG* | Invitrogen |
| *E. coli* ET12567/pUZ8002 | A methylation deficient ($\Delta dcm\Delta dam$) strain of *E. coli* containing the driver plasmid pUZ8002 for conjugation to *Streptomyces venezuelae*. | John Innes Centre<br>(http://streptomyces.org.uk) |
| *E. coli* BL21 + pETDuet-*mtrA* | Non-T7 *E. coli* expression strain used for purification of 6x His-MtrA<br>*fhuA2 [lon] ompT gal [dcm] $\Delta hsdS$* | New England Biolabs |
| *S. venezuelae* NRRL B-65542 | Wild-type *S. venezuelae* NRRL B-65542 | John Innes Centre<br>(http://streptomyces.org.uk) |
| *S. venezuelae* $\Delta mtrA$ | *S. venezuelae* NRRL B-65542 with an in-frame deletion of *mtrA* generated using pCRISPomyces-2 | This work |
| *S. venezuelae* $\Delta mtrA$ + *mtrA* | *S. venezuelae* $\Delta mtrA$ complemented *in trans* by integration of pSS170 containing *mtrA* under the control of the native promoter | This work |
| *S. venezuelae* $\Delta mtrA$ + *PermE\*-bldM-whiI* | *S. venezuelae* $\Delta mtrA$ containing integrated pIJ10257 with an artificial *bldM-whiI* operon under the control of the high-level constitutive *ermE\** promoter. | This work |

**Table 2.**
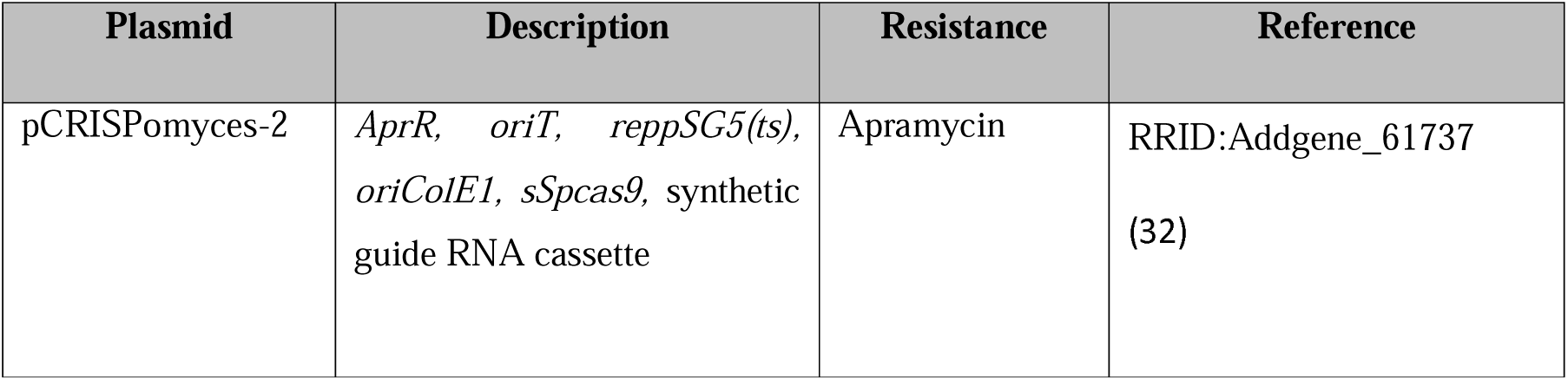
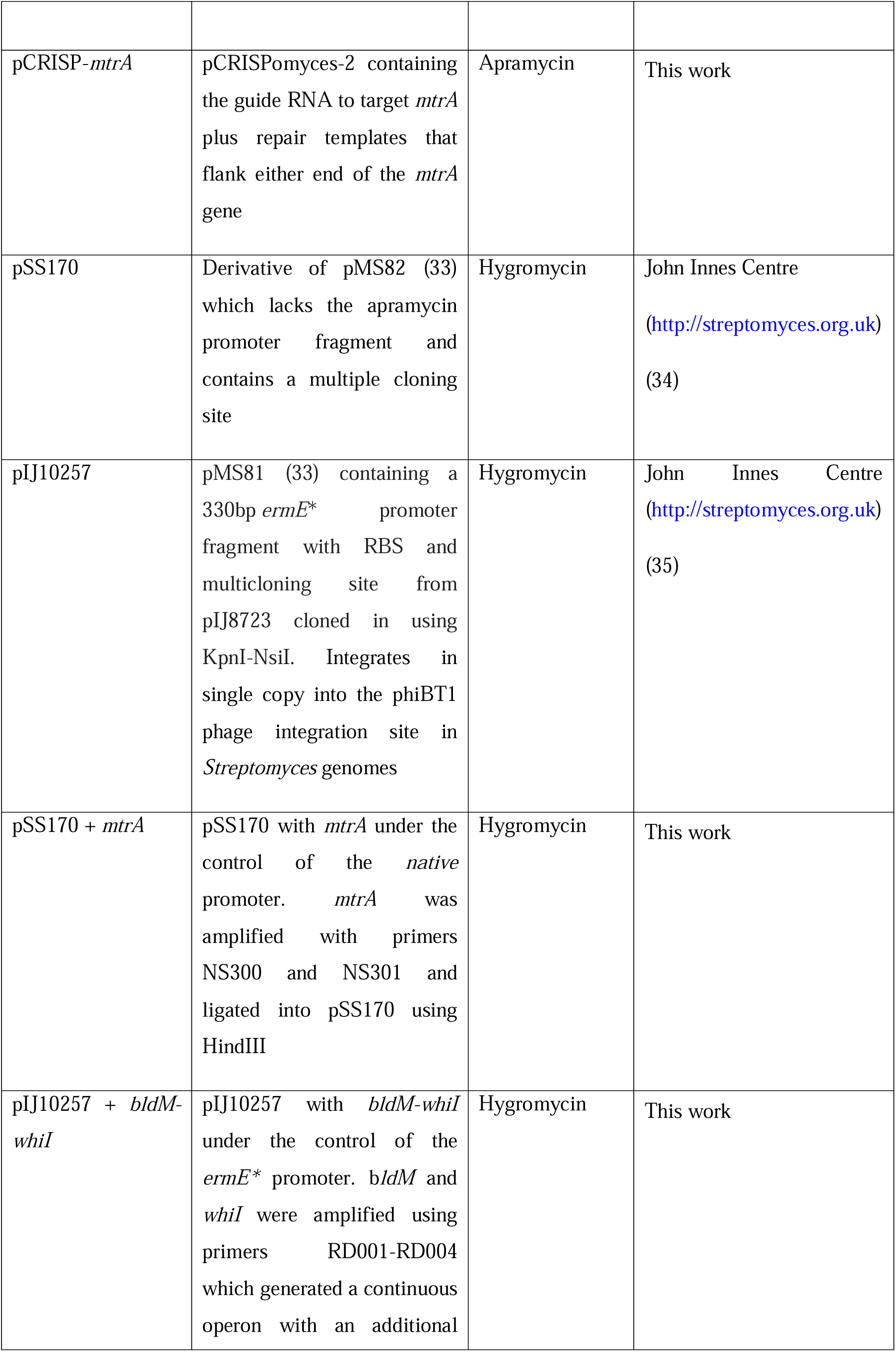
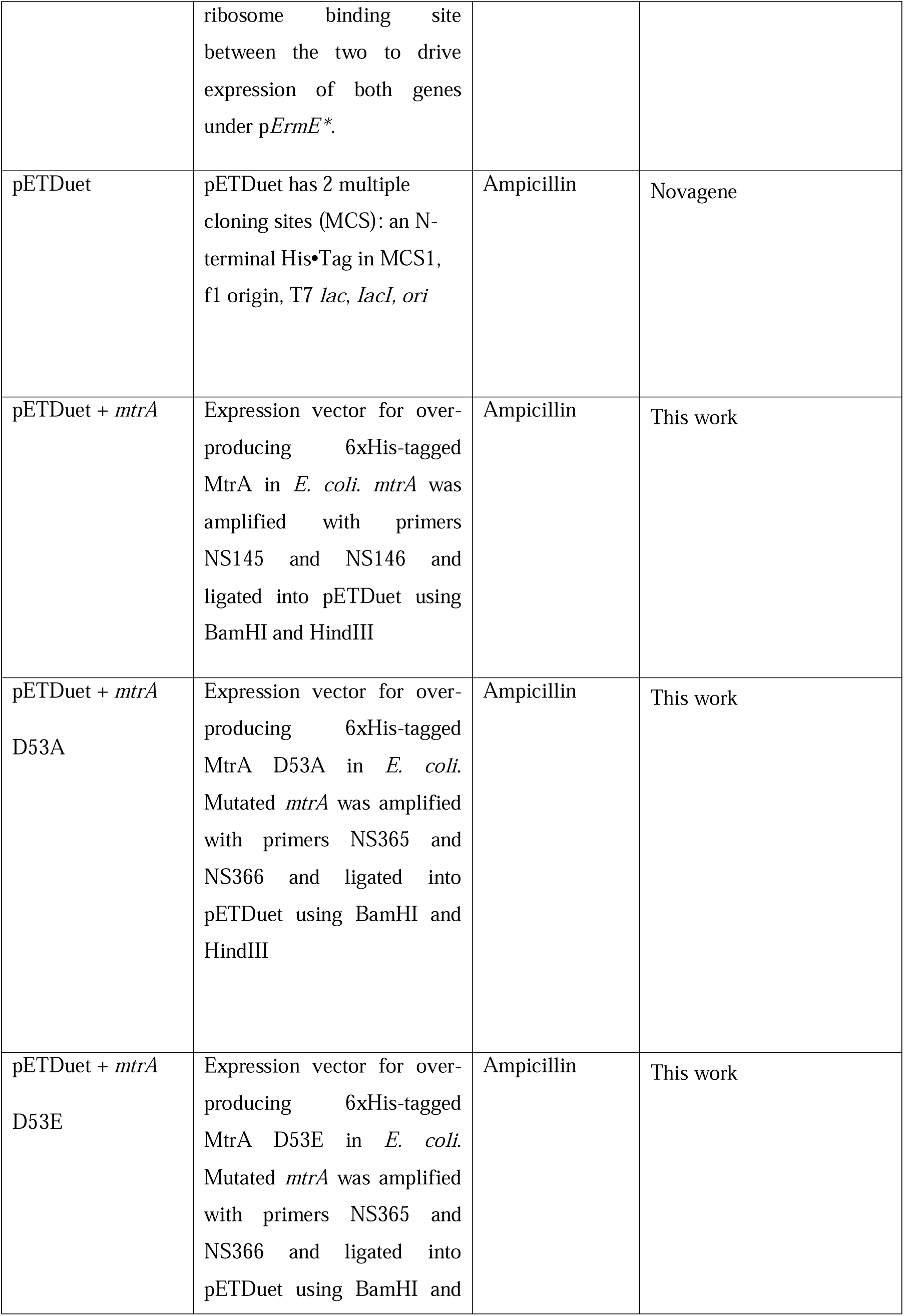
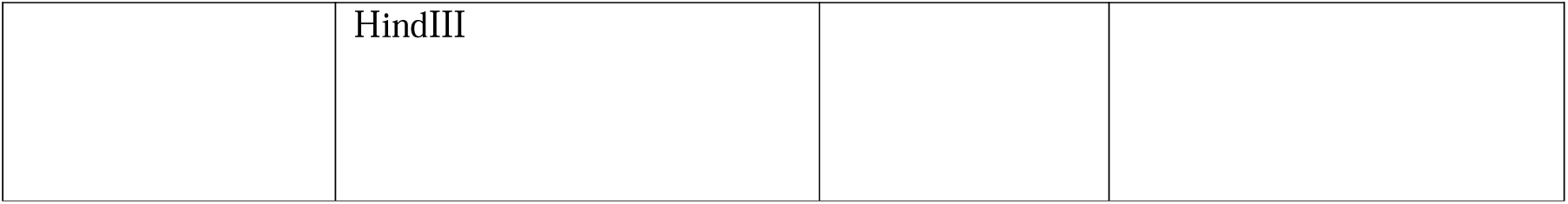
Plasmids used in this work.

| Plasmid | Description | Resistance | Reference |
| --- | --- | --- | --- |
| pCRISPomyces-2 | *AprR*, *oriT*, *reppSG5(ts)*, *oriColEI*, *sSpcas9*, synthetic guide RNA cassette | Apramycin | RRID:Addgene_61737<br>(32) |
| pCRISP-*mtrA* | pCRISPomyces-2 containing the guide RNA to target *mtrA* plus repair templates that flank either end of the *mtrA* gene | Apramycin | This work |
| pSS170 | Derivative of pMS82 (33) which lacks the apramycin promoter fragment and contains a multiple cloning site | Hygromycin | John Innes Centre<br>http://streptomyces.org.uk<br>(34) |
| pIJ10257 | pMS81 (33) containing a 330bp *ermE* * promoter fragment with RBS and multicloning site from pIJ8723 cloned in using KpnI-NsiI. Integrates in single copy into the phiBT1 phage integration site in *Streptomyces* genomes | Hygromycin | John Innes Centre<br>http://streptomyces.org.uk<br>(35) |
| pSS170 + *mtrA* | pSS170 with *mtrA* under the control of the *native* promoter. *mtrA* was amplified with primers NS300 and NS301 and ligated into pSS170 using HindIII | Hygromycin | This work |
| pIJ10257 + *bldM-whiI* | pIJ10257 with *bldM-whiI* under the control of the *ermE* * promoter. *bldM* and *whiI* were amplified using primers RD001-RD004 which generated a continuous operon with an additional ribosome binding site between the two to drive expression of both genes under p *ErmE* *. | Hygromycin | This work |
| pETDuet | pETDuet has 2 multiple cloning sites (MCS): an N-terminal His•Tag in MCS1, f1 origin, T7 *lac*, *lacI*, *ori* | Ampicillin | Novagene |
| pETDuet + *mtrA* | Expression vector for over-producing 6xHis-tagged MtrA in *E. coli*. *mtrA* was amplified with primers NS145 and NS146 and ligated into pETDuet using BamHI and HindIII | Ampicillin | This work |
| pETDuet + *mtrA* D53A | Expression vector for over-producing 6xHis-tagged MtrA D53A in *E. coli*. Mutated *mtrA* was amplified with primers NS365 and NS366 and ligated into pETDuet using BamHI and HindIII | Ampicillin | This work |
| pETDuet + *mtrA* D53E | Expression vector for over-producing 6xHis-tagged MtrA D53E in *E. coli*. Mutated *mtrA* was amplified with primers NS365 and NS366 and ligated into pETDuet using BamHI and | Ampicillin | This work |
|  | HindIII |  |  |

### Growth media and conditions

The growth media used in this work are listed in Table 3. Recipes are also available at https://actinobase.org/index.php?title=Media_recipes. The final concentrations of antibiotics used for selection of plasmid carriage were 50 μg/ml of apramycin or 50 μg/ml hygromycin. Spores were pregerminated at 50°C for 10 minutes in 2x Yeast Tryptone (2xYT) prior to conjugation with *E. coli.* After conjugation, 25 μg/ml of nalidixic acid was used to kill the *E. coli* cells alongside the relevant antibiotic to select for *S. venezuelae* exconjugants.

**Table 3.**
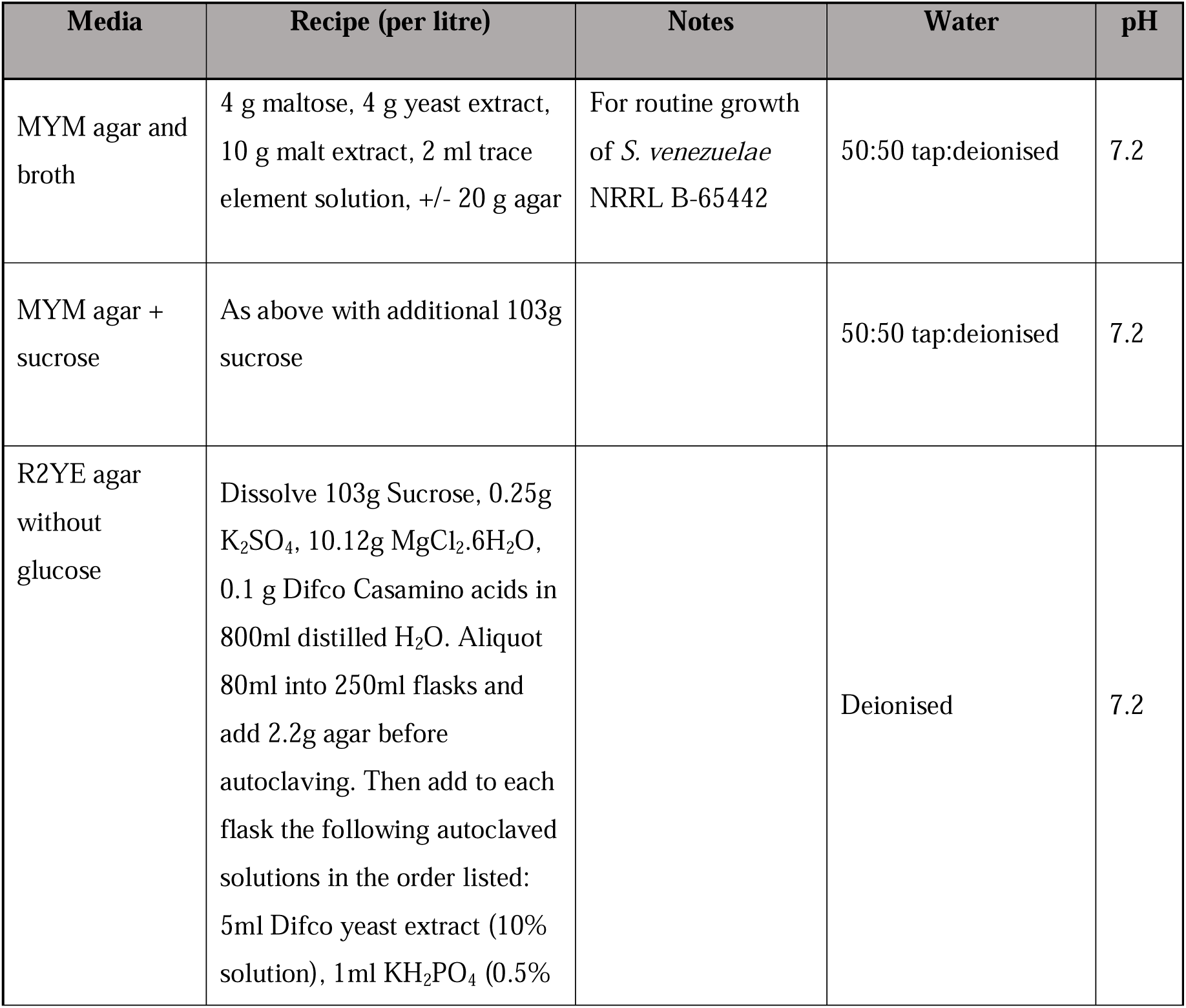
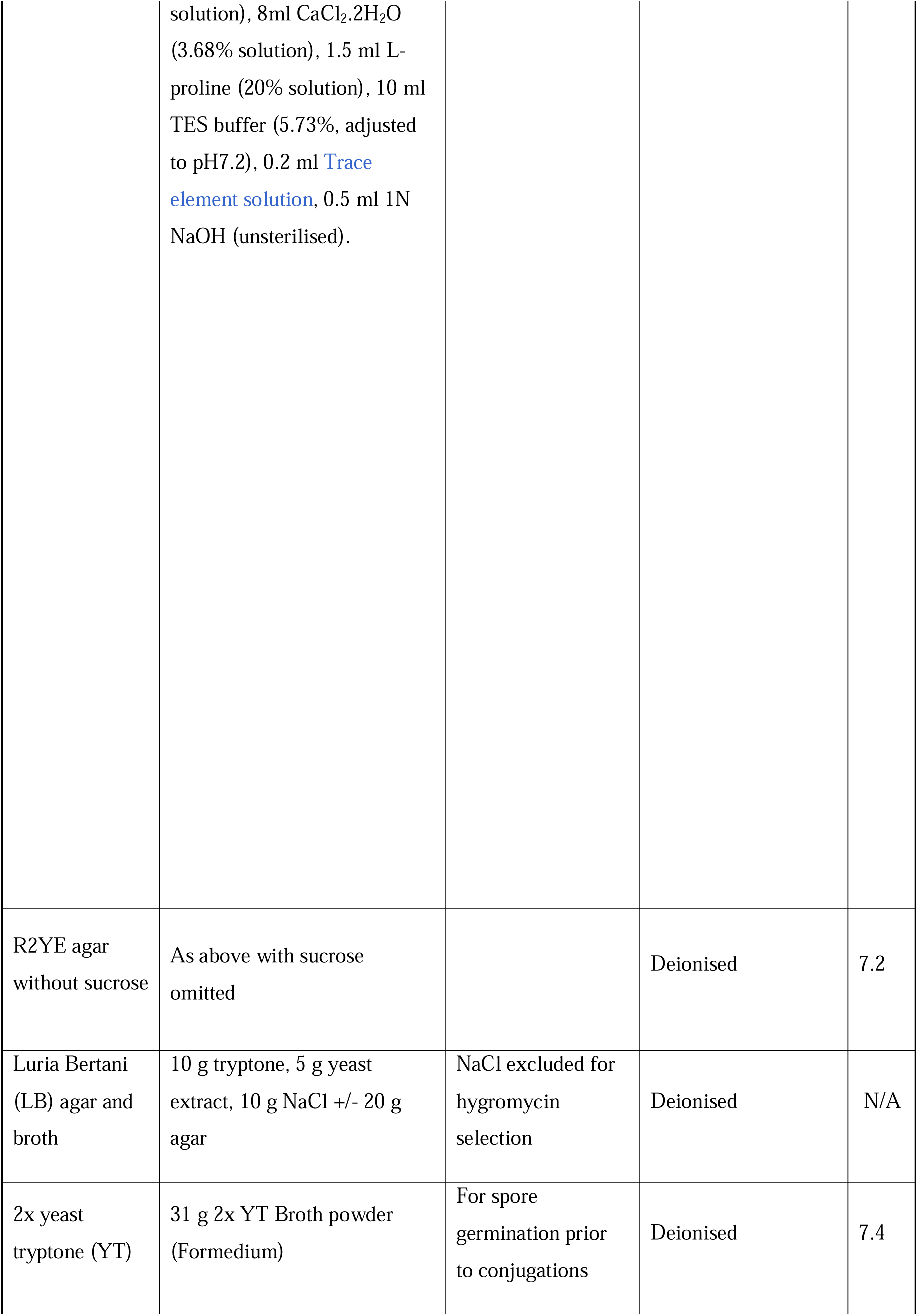
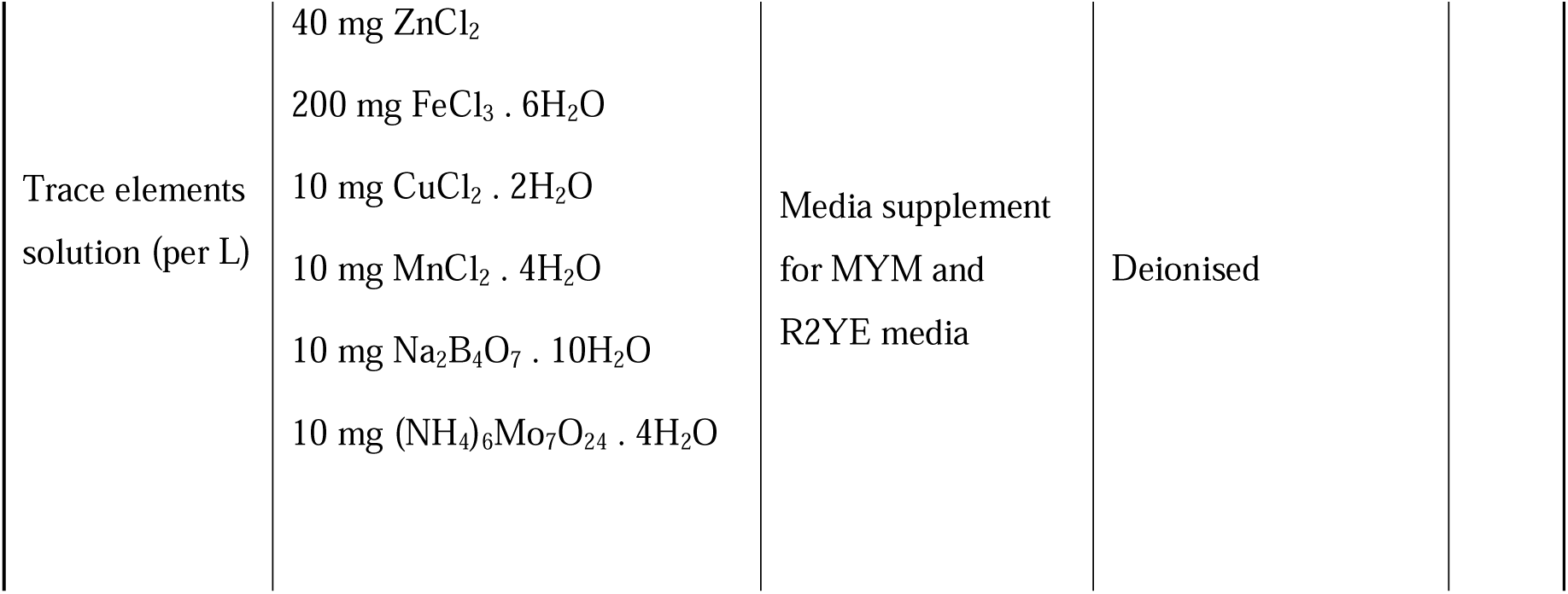
Growth media and supplements used for bacterial culturing.

| Media | Recipe (per litre) | Notes | Water | pH |
| --- | --- | --- | --- | --- |
| MYM agar and broth | 4 g maltose, 4 g yeast extract, 10 g malt extract, 2 ml trace element solution, +/- 20 g agar | For routine growth of *S. venezuelae* NRRL B-65442 | 50:50 tap:deionised | 7.2 |
| MYM agar + sucrose | As above with additional 103g sucrose |  | 50:50 tap:deionised | 7.2 |
| R2YE agar without glucose | Dissolve 103g Sucrose, 0.25g K<sub>2</sub>SO<sub>4</sub>, 10.12g MgCl<sub>2</sub>.6H<sub>2</sub>O, 0.1 g Difco Casamino acids in 800ml distilled H<sub>2</sub>O. Aliquot 80ml into 250ml flasks and add 2.2g agar before autoclaving. Then add to each flask the following autoclaved solutions in the order listed: 5ml Difco yeast extract (10% solution), 1 ml KH<sub>2</sub>PO<sub>4</sub> (0.5% |  | Deionised | 7.2 |
|  | solution), 8ml CaCl<sub>2</sub>.2H<sub>2</sub>O (3.68% solution), 1.5 ml L-proline (20% solution), 10 ml TES buffer (5.73%, adjusted to pH7.2), 0.2 ml Trace element solution, 0.5 ml 1N NaOH (unsterilised). |  |  |  |
| R2YE agar without sucrose | As above with sucrose omitted |  | Deionised | 7.2 |
| Luria Bertani (LB) agar and broth | 10 g tryptone, 5 g yeast extract, 10 g NaCl +/- 20 g agar | NaCl excluded for hygromycin selection | Deionised | N/A |
| 2x yeast tryptone (YT) | 31 g 2x YT Broth powder (Formedium) | For spore germination prior to conjugations | Deionised | 7.4 |

|  |  |  |  |
| --- | --- | --- | --- |
| Trace elements<br>solution (per L) | 40 mg ZnCl<sub>2</sub> | Media supplement<br>for MYM and<br>R2YE media | Deionised |
|  | 200 mg FeCl<sub>3</sub> · 6H<sub>2</sub>O |  |  |
|  | 10 mg CuCl<sub>2</sub> · 2H<sub>2</sub>O |  |  |
|  | 10 mg MnCl<sub>2</sub> · 4H<sub>2</sub>O |  |  |
|  | 10 mg Na<sub>2</sub>B<sub>4</sub>O<sub>7</sub> · 10H<sub>2</sub>O |  |  |
|  | 10 mg (NH<sub>4</sub>)<sub>6</sub>Mo<sub>7</sub>O<sub>24</sub> · 4H<sub>2</sub>O |  |  |

### Deletion of *mtrA* using Cas9-mediated genome editing

To make an in-frame deletion of the *mtrA* gene a guide RNA was designed and then generated by annealing primers NS308 and NS309 and cloned into the pCRISPomyces-2 plasmid using Golden Gate (32). The vector containing the guide RNA was then linearised using XbaI to clone in the repair templates (generated using primers NS304-307) using Gibson Assembly as described previously. The resulting vector was named pCRISP-*mtrA*. A detailed protocol for the use of pCRISPomyces-2 is available at: https://actinobase.org/index.php?title=CRISPR/Cas9-mediated_genome_editing.

### Purification of MtrA

6xHis-tagged MtrA was over-produced in *E. coli* BL21 using pETDuet. Cultures were grown in 2L LB, shaking at 37 °C, 200 rpm for 4 hours, then induced with 100 µM IPTG and incubated at 18 °C overnight. Cells were harvested by centrifugation and lysed by sonication on ice in 20 mM Tris pH 8, 75 nM NaCl, 0.1% v/v Triton-X, 10 µg/ml lysozyme, EDTA-free protease inhibitors. The soluble fraction was harvested by centrifugation at 18000 rpm for 20 minutes; this was then applied directly to a nickel affinity chromatography (HiTrap) column on an AKTApure in a refrigerated cabinet. The column had been equilibrated with wash buffer (80 mM Tris pH 8, 200 mM NaCl, 1 mM TCEP, 20 mM imidazole). After further rounds of column washing to remove non-specifically bound proteins, MtrA was eluted from the column using wash buffer + 350 mM imidazole and fractions were analysed by SDS-PAGE to confirm the presence of the protein.

### Reuseable DNA Capture with Surface Plasmon Resonance (ReDCaT SPR)

Oligonucleotides (Table S2) were designed against key developmental promoters (Table 5) and annealed in HBS-EP+ Buffer to make double stranded probes. DNA probes were annealed to a streptavidin chip prepared with ReDCaT linker DNA using the Biocore 8K+ SPR system in HBS-EP+ Buffer and water. Oligos were flowed over the chip at 10 ul/min for a 60 second capture, followed by freshly purified MtrA protein to analyse DNA binding activity. Binding was recorded at both early and late time points, after 60 seconds contact followed by 60 seconds dissociation. The chip was regenerated using a 60 second wash with 1M NaCl/50 mM NaOH reagent to remove protein and DNA probes for the next cycle. Binding responses were normalised for the molecular weight of both DNA and protein.

### Tandem-mass-tag (TMT) proteomics

*S. venezuelae* colonies were grown on cellophane covered MYM or R2YE – glucose agar plates as triplicate spots from 5 µl spores, all in duplicate. After 5 days at 30 °C the mycelium was scraped into a 15 ml Falcon tube and resuspended in 10 ml cell lysis buffer [50 mM TEAB buffer pH 8.0, 150 mM NaCl, 2% SDS, EDTA-free protease inhibitor, PhosSTOP phosphatase inhibitor (Sigma Aldrich)]. The suspension was disrupted via French press three times before being boiled for 10 min. Samples were sonicated at 50 kHz four times for 20 seconds per cycle and then pelleted at 3,220 x g for 30 min. Protein concentration was determined using the BCA assay and 1 mg of protein from each sample transferred to a fresh 15 ml Falcon tube. Four volumes of methanol were added, the sample was vortexed, then 1 volume Chloroform and 3 volumes of dH_2_O. Samples were centrifuged at 4000 xg for 10 minutes and the upper layer disgarded, leaving the interphase behind. Another 4 volumes of methanol were added to the bottom phase and then vortexed, followed by thorough vortexing and centrifugation at 4000 xg for 20 minutes. As much solvent as possible was removed without disturbing the pellet and 3ml acetone added. This was incubated for 10 minutes on the bench, then transferred to a mircocentrifuge tube for centrifugation at 13000 rpm for 20 minutes. The acetone was removed and the pellet air dried before continuing.

The protein pellets were resuspended in 250 µl of 2.5% sodium deoxycholate (SDC; Merck) in 0.2 M EPPS-buffer (Merck), pH 8.5, and vortexed under heating. Protein concentration was estimated using a BCA assay. For the analysis, approx. 100 µg of protein per sample was used. The proteins were treated with DTT (10 mM, 60°C, 45 min) and iodoacetamide (30 mM, 45 min) to reduce and alkylate cysteine residues. One µg of Trypsin (Sequencing Grade, Promega) was added per sample and the samples were incubated at 40°C for about 18 h. After the digest, TMT labelling was performed using 12 channels from a TMT™16plex kit (Lot ZD386952, ThermoFisher Scientific) according to the manufacturer’s instructions. Samples were assigned to the TMT channels in alternating order to avoid channel leakage according to Brenes *et al*. between different conditions with channels 126, 130N, 130C, 131N omitted (36). After 2 h incubation, aliquots of 2 µl from each sample were combined in 600 µl 0.5% TFA, desalted, and analysed on the mass spectrometer (same method as for TMT, but without RTS, see below) to test labelling efficiency and estimate total sample abundances. The main sample aliquots were quenched by adding 8 µl of 5% hydroxylamine and then combined to roughly level abundances according to the test and desalted using a C18 Sep-Pak cartridge (200 mg, Waters). The eluted peptides were dissolved in 500 µl of 25 mM NH_4_HCO_3_ and fractionated by high pH reversed phase HPLC. Using an ACQUITY Arc Bio System (Waters), the samples were loaded to an XBridge® 5 µm BEH C18 130 Å column (250 x 4.6 mm, Waters). Fractionation was performed with the following gradient of solvents A (water), B (acetonitrile), and C (25 mM NH_4_HCO_3_ in water, pH 8.5) at a flow rate of 1 ml min^-1^: solvent C was kept at 10% throughout the gradient; solvent B: 0-5 min: 5%, 5-10 min: 5-10%, 10-80 min: 10-45%, 80-90 min: 45-80%, followed by 5 min at 80% B and re-equilibration to 5% for 25 min. Fractions of 1 ml were collected and concatenated by combining distant fractions of similar peptide concentration to produce 23 final fractions for MS analysis. Aliquots were analysed by nanoLC-MS/MS on an Orbitrap Eclipse™ Tribrid™ mass spectrometer equipped with a FAIMS Pro Duo interface and coupled to a Vanquish™ Neo UHPLC System (Thermo Fisher Scientific, Hemel Hempstead, UK). The samples were loaded onto a trap cartridge (PepMap™ Neo Trap Cartridge, C18, 5um, 0.3x5mm, Thermo) with 10 µl of 0.1% TFA at 15 µl min^-1^. The trap column was then switched in-line with the analytical column (Aurora Frontier TS, 60 cm nanoflow UHPLC column, ID 75 µm, reversed phase C18, 1.7 µm, 120 Å; IonOpticks, Fitzroy, Australia) for separation at 60°C using the following gradient of solvents A (water, 0.1% formic acid) and B (80% acetonitrile, 0.1% formic acid) at a flow rate of 0.25 µl min^-1^: 0-7 min increase B from 1% to 7% (curve 4), 7-107 min increase B to 46% (or 52% depending on fraction); 107-112 min linear increase B to 99 %; keeping at 99% B for 3 min and re-equilibration to 1% B. Data were acquired with the following parameters in positive ion mode: MS1/OT: resolution 120k, profile mode, mass range *m/z* 400-1600, AGC target 4e^5^, max inject time 50 ms, FAIMS device set to three compensation voltages (-35V, -50V, -65V) for 1 s each; MS2/IT: for each CV, data dependent analysis with the following parameters: 1 s cycle time Rapid mode, centroid mode, quadrupole isolation window 0.7 Da, charge states 2-5, threshold 1.9e^4^, CID CE=30, AGC target 1e4, max. inject time 50 ms, dynamic exclusion 1 count for 30 s, mass tolerance ± 7 ppm; MS3 synchronous precursor selection (SPS): 10 SPS precursors, isolation window 0.7 Da, HCD fragmentation with CE=50, Orbitrap Turbo TMT and TMTpro resolution 30k, AGC target 200%, max inject time 100 ms, Real Time Search (RTS): protein database *Streptomyces venezuelae,* enzyme trypsin, 1 missed cleavage, oxidation (M) as variable, carbamidomethyl (C) and TMTpro as fixed modifications, precursor tolerance 10 ppm, match parameters: Xcorr = 1.4, dCn = 0.1.

The mass spectrometry raw data were processed and quantified in Proteome Discoverer 3.2 (Thermo); all mentioned tools of the following workflow are nodes of the proprietary Proteome Discoverer (PD) software. The *S. venezuelae database* was imported into PD adding a reversed sequence database for decoy searches; a database for common contaminants (maxquant.org, 245 entries) was also included. The database search was performed using the incorporated search engines CHIMERYS (MSAID, Munich, Germany) and Comet (37). The processing workflow included recalibration (RC) of MS1 spectra, reporter ion quantification by most confident centroid (20 ppm) and a search on the imported *S. venezuelae* database. The TopN Peak Filter was applied with 20 peaks per 100 Da. For CHIMERYS, the inferys_4.7.0._fragmentation prediction model was used with fragment tolerance of 0.3 Da, enzyme trypsin with 1 missed cleavage, variable modification oxidation (M), fixed modifications carbamidomethyl (C) and TMTpro on N-terminus and K. For Comet the version 2019.01 rev.0 parameter file was used with default settings except precursor tolerance set to 5 ppm and trypsin missed cleavages set to 1. Modifications were the same as for CHIMERYS. The consensus workflow included the following parameters: use of intensity-based abundance, normalisation on total peptide abundances, protein abundance-based ratio calculation, only unique peptides (protein groups) for quantification, TMT channel correction values applied (Lot ZD386952), co-isolation/SPS matches/CHIMERYS Coefficient thresholds 50%/65%, 0.8; missing values imputation by low abundance resampling, hypothesis testing by t-test (background based), adjusted p-value calculation by the Benjamini-Hochberg procedure. The results were filtered for strict FDR confidence (0.01) and Master proteins and were exported into a Microsoft Excel table including, among others, results for normalised and un-normalised abundances, ratios for the specified conditions calculated from the normalised abundances, the corresponding p-values and adjusted p-values, number of unique peptides, q-values, PEP-values, identification scores from both search engines.

### Ectoine extraction and quantification

Strains were grown in lawns for 3 days (MYM) or 7 days (R2YE) and small plugs of equal sizes were taken for extraction (n=3). Plugs were freeze-thawed to assist with cell lysis before being submerged in ice-cold 50:50 water:methanol (400 µl). Samples were sonicated on ice until lysed (15 seconds on, 15 seconds off, for 3 minutes) and incubated on ice for a further 20 minutes with occasional vortexing. Samples were then centrifuged at 13000 rpm for 10 minutes at 4 °C to pellet the cell debris. Supernatant was collected and dried in a GeneVac evaporator. Samples were resuspended in 90% Acetonitrile with 10% dH_2_O + 0.1% formic acid and diluted 1:10 for analysis. The samples were analysed by liquid chromatography coupled with electrospray ionisation tandem mass spectrometry (LC-ESI-MS/MS) on TQ-Absolute triple quadrupole mass spectrometer (Waters) operated in MRM mode. ESI-MS/MS analysis was performed in positive ion mode using a source with a capillary voltage of 3 kV, 600 °C desolvation temperature, 900 l/h desolvation gas, 150 l/h cone gas, and 7 bar nebulizer pressure. MRM transitions for ectoine standard (Sigma-Aldrich) in positive ESI mode (Table 4) were generated using IntelliStart software. Sample of ectoine (10 μM) was introduced at 10 μl/min combined with a flow from the UPLC pump typical of an LC run. Liquid chromatography was achieved on an Accucore 150-Amide-HILIC (100 x 2.1 mm, 2.6 µm) column equipped with a column guard. Ectoine was eluted using mobile phase A: water containing 0.1% formic acid and mobile phase B: acetonitrile using the following multistep gradient at a flow rate 400 μl/min: 0 min: 95% B; 1 min 95% B; 10 min: 60% B; 11 min: 60% B; 12 min: 95% B; 20 min: 95% B. Ectoine standard (10 μM) was injected (1 μl) to determine retention time. Once LC retention time of ectoine has been established, the mass transitions were collected in time-windows centred on the relevant peak. MassLynx software (Waters) was used to collect, to analyse and to process data. Limit of detection for ectoine was determined to be 10 fmol on column using a serial dilution. Transition 143 → 97 was used for quantification.

**Table 4:** MRM transitions of ectoine.

| Standard | MRM transitions | Cone [V] | Collision Energy [eV] | Dwell [s] |
| --- | --- | --- | --- | --- |
| Ectoine | 143 → 68 | 2.00 | 22.00 | 1.497 |
| | 143 → 97 | 2.00 | 16.00 | 1.497 |

### Cryo-Scanning Electron Microscopy

Samples were prepared for cryo scanning electron microscopy (cryo-SEM) by streaking for single colonies on the appropriate media as described above and incubated at 30°C for the indicated number of days. Samples were mounted on an aluminium stub using Tissue-Tek^R^ (BDH Laboratory Supplies, Poole, England). The stub was immediately plunged into liquid nitrogen slush (approximately −210 °C) to cryo-preserve the material. The frozen sample was then transferred to the cryo-stage of an ALTO 2500 cryo-transfer system (Gatan, Oxford, England) attached to an FEI Nova NanoSEM 450 (FEI, Eindhoven, The Netherlands).

Surface frost was sublimated at −95 °C for 4 minutes, after which the sample was sputter-coated with platinum for 130 seconds at 10 mA, at a temperature below −110 °C. The sample was then transferred to the main chamber of the microscope, where the cryo-stage was maintained at approximately −125 °C. Imaging was performed at 3 kV, and digital TIFF files were recorded.

## Results and Discussion

### Deletion of *S. venezuelae mtrA* prevents sporulation on R2YE agar

Previous studies have shown that deletion of *mtrA* results in a conditional bald phenotype in *S. coelicolor* M145, *S. lividans* and *S. venezuelae* ISP5230. This means the Δ*mtrA* mutants can sporulate on some growth media but not others, for example Δ*mtrA* mutants of all three species fail to sporulate on R2YE and R5 agar (22,29). To test whether *mtrA* deletion blocks sporulation in *S. venezuelae* NRRL B-65442 we deleted the *mtrA* gene using pCRISPomyces-2 (32) and screened the Δ*mtrA* mutant for defects in growth, development and antibiotic production on MYM agar, a sporulation medium, and on R2YE agar. Glucose was omitted from the R2YE agar recipe because it prevents the wild-type strain from sporulating. The results show that the Δ*mtrA* mutant sporulates normally on MYM agar but fails to develop aerial hyphae or spores on R2YE agar, indicating that *S. venezuelae* NRRL B-65542 Δ*mtrA* is conditionally bald (Figure 1A). This was confirmed using scanning electron microscopy (Figure 1B).

**Figure 1.**
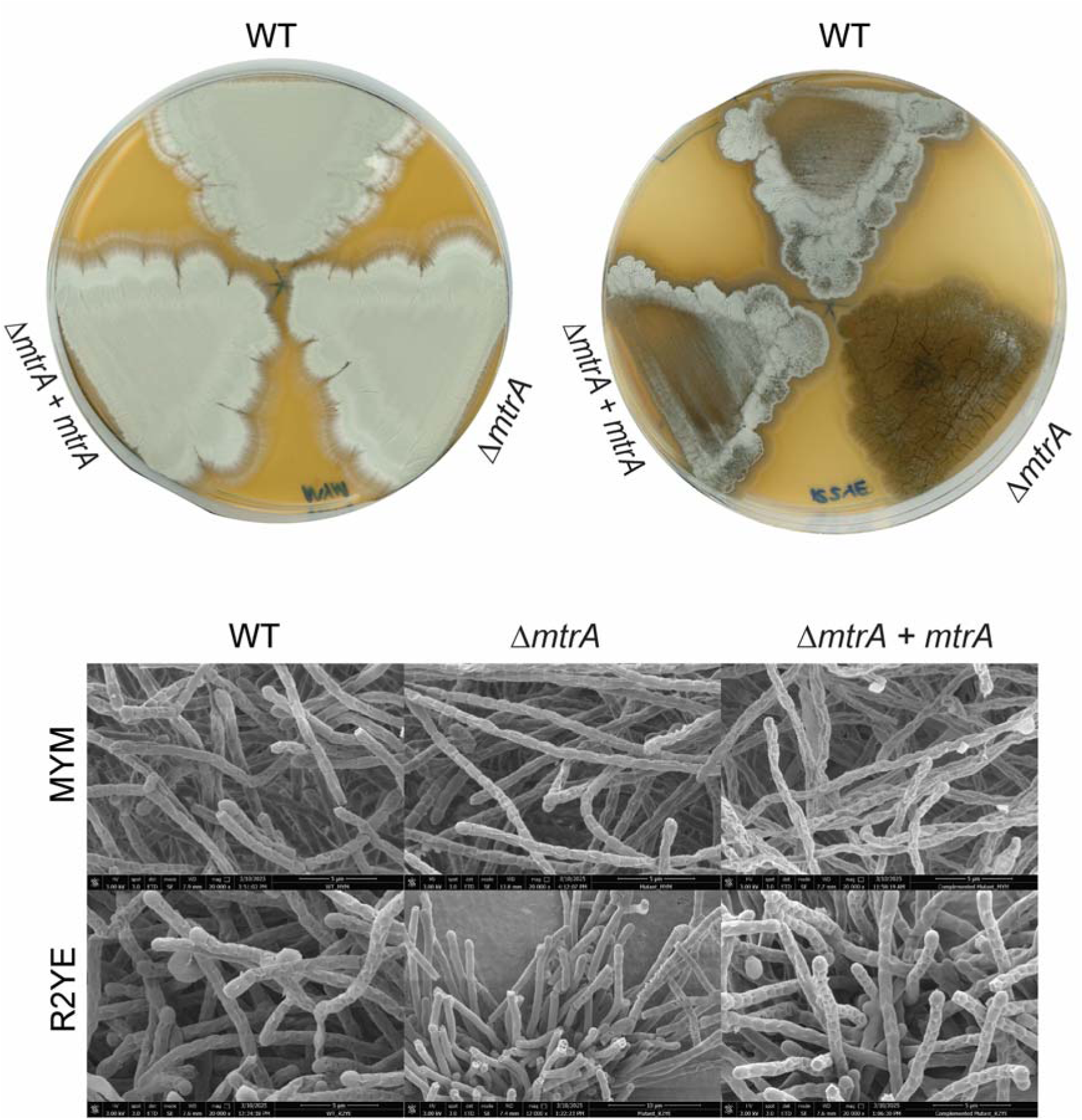
*S. venezuelae* Δ*mtrA* is conditionally bald and cannot sporulate on R2YE agar. **A.** Seven day old cultures of *S. venezuelae* wild-type (WT), Δ*mtrA* and the Δ*mtrA* strain complemented *in trans* grown on MYM agar (left) and R2YE agar without glucose (right). Aerial mycelium is white and the spore chains are green due to production of the green WhiE spore-pigment. **B.** Scanning electron micrographs of the same strains grown under the same conditions.

### Identification of MtrA binding sites upstream of genes required for hyphal growth, DNA replication and sporulation

We analysed existing *S. venezuelae* MtrA ChIP-seq datasets (1) to identify developmental genes with ChIP-seq peaks in their promoters or coding sequences, which indicate MtrA binding *in vivo*. MtrA binding was detected upstream of the developmental genes *adpA*, *bldM, filP*, *mtrA*, *ssgB* and *whiI*, which are required for aerial hyphae formation and sporulation, and upstream of the *dnaA* gene which is essential for DNA replication (Figure S1). These genes were chosen for further investigation because MtrA-dependent changes in their expression could be responsible for the conditional bald phenotype of the *S. venezuelae* Δ*mtrA* mutant. To pinpoint the precise MtrA binding motifs at these promoters, we used <u>Re</u>usable <u>D</u>NA <u>Ca</u>pture <u>T</u>echnology <u>S</u>urface <u>P</u>lasmon <u>R</u>esonance (ReDCaT SPR). This technique involves designing double stranded oligonucleotide probes that tile across a target promoter to narrow down the MtrA binding site (38). Truncation and mutation of positive probes can then be used to identify the DNA sequence required for MtrA binding. The results show that all of these promoters contain recognisable MtrA binding sites that are bound by purified MtrA *in vitro*, with some promoters (*adpA*, *bldM* and *dnaA*) containing two or more MtrA binding sites (Table 5).

**Table 5.**
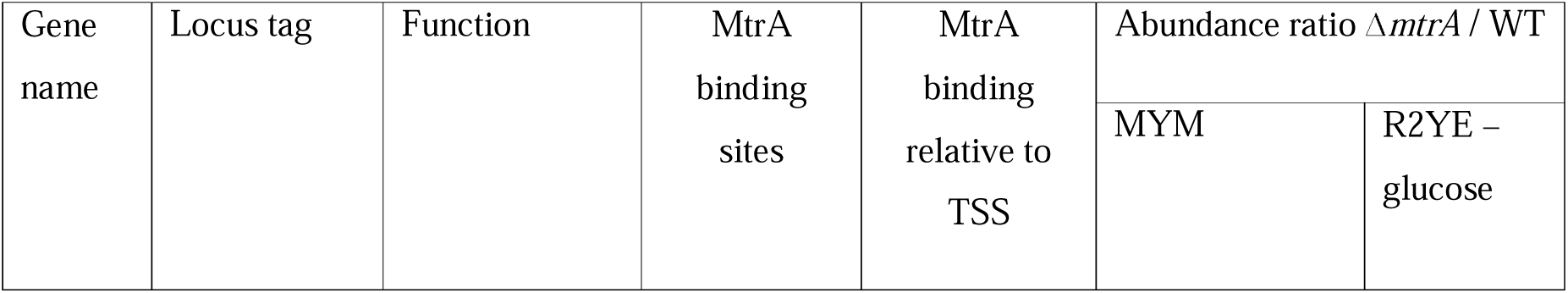
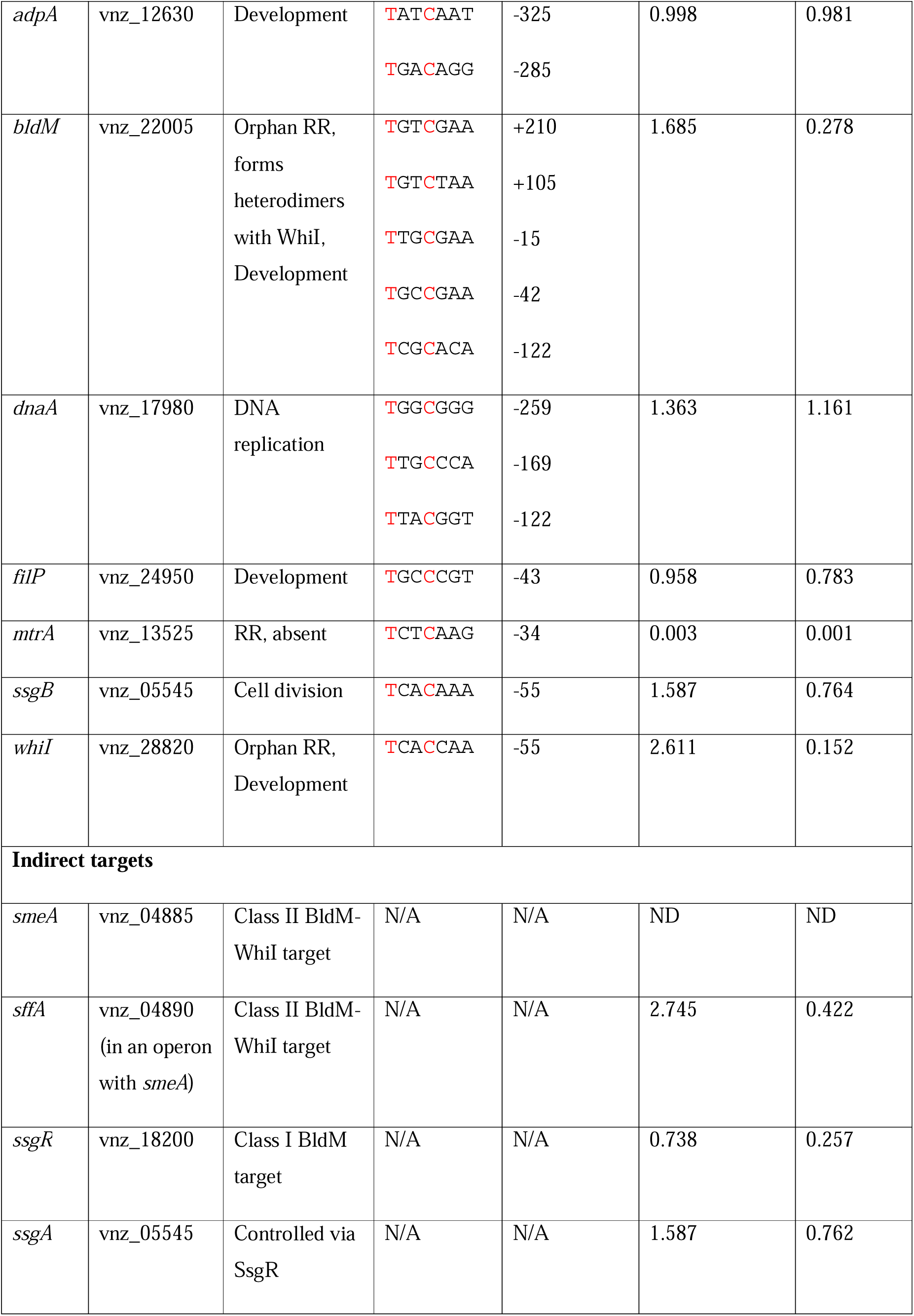
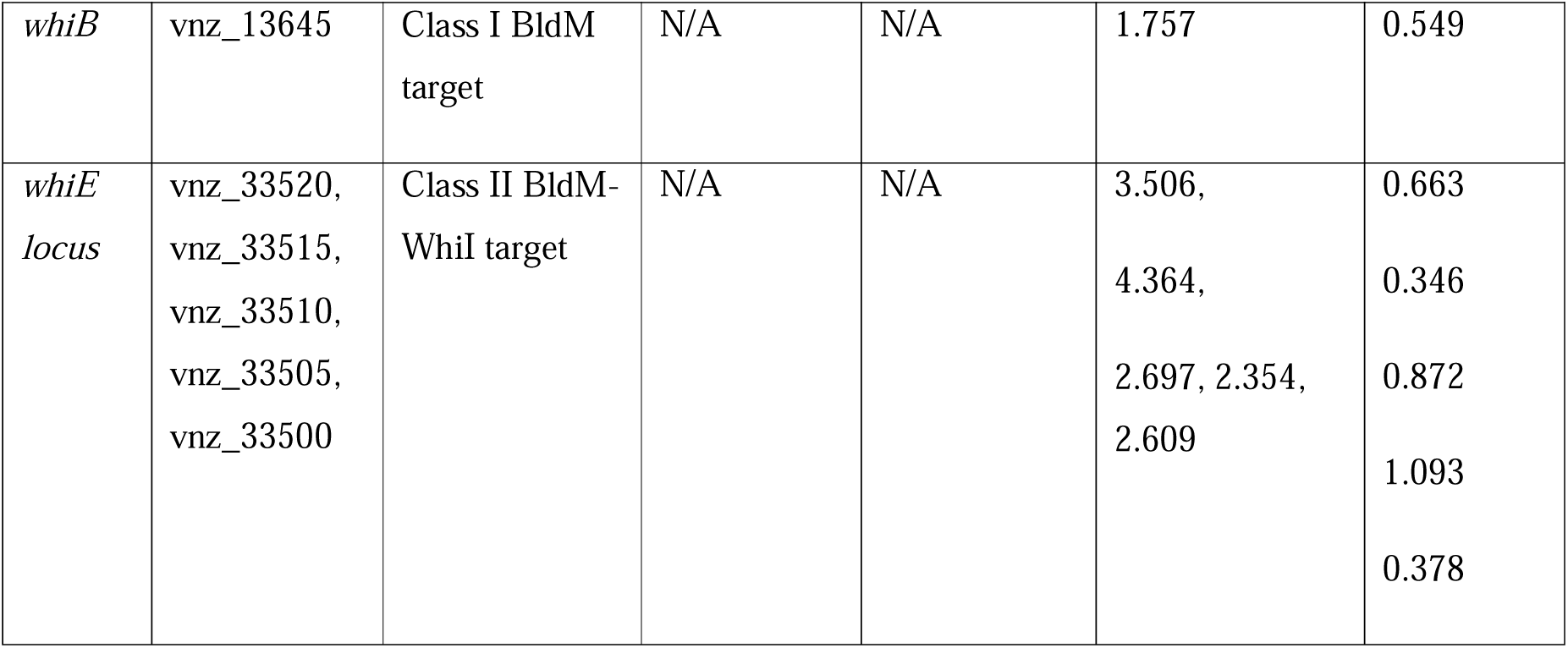
MtrA target promoters upstream of *S. venezuelae* developmental genes. The MtrA binding sites were identified using ReDCaT SPR and positions are given relative to the transcript start sites (TSS) of the downstream gene. Tandem mass tag (TMT) proteomics was used to measure the levels of each gene product after growth on either MYM or R2YE agar. The MtrA binding sites and TSS locations are given relative to the translational start codon of each gene. Note that MtrA is missing from the Δ*mtrA* mutant. Genes can be viewed using the StrepDB genome browser at http://streptomyces.org.uk, select “vnz chr” from the drop down Org & Mol list and search by gene number (e.g. vnz_12630). In all cases an abundance ratio Δ*mtrA* / WT of >2.0 or <0.7 has a p-value of <0.05 all others represent no significant change between Δ*mtrA* and WT

### Defining the MtrA consensus binding sequence

Alignment of the 14 experimentally verified MtrA binding sites shown in Table 1 results in a seven base pair consensus sequence (Figure 2). This seven base pair consensus site sits at the 3’ edge of the *mtrA* promoter probe, so we sequentially truncated the probe until binding was abolished. Results show that any truncation of the 3’ edge of the probe affects the ability of MtrA to bind, and removal of the 15bp containing the identified consensus sequence abolishes binding. To further confirm this sequence in the *mtrA* promoter as the MtrA binding site, the 7bp sequence was scrambled within the DNA probe, and this significantly reduced the ability of MtrA to bind (Figure 2A). Notably, only two nucleotides in the consensus DNA binding site, T at position 1 and C at position 4, are conserved across all identified hits, and mutation of either of these significantly reduces MtrA binding (Figure 2B).

**Figure 2:**
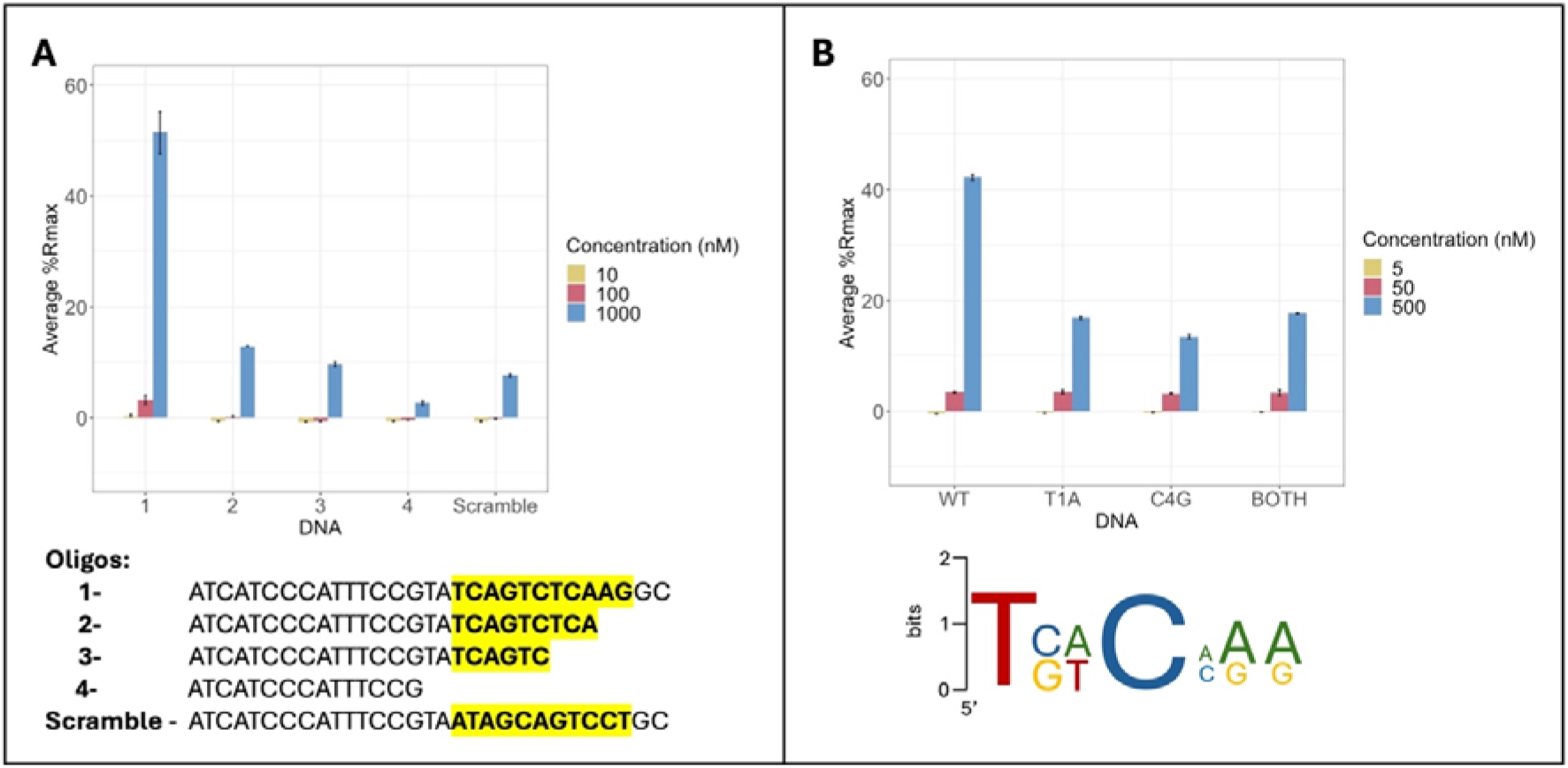
Consensus MtrA binding site. **A:** Alignment of the probes bound by MtrA *in vitro* revealed a 7bp consensus sequence. This binding site is at the 3’ end of the *mtrA* promoter. Removal of this site (highlighted) by sequential truncation of the *mtrA* promoter probe abolished DNA binding. Similarly, scrambling the 7bp binding site within this oligo also significantly reduces DNA binding by purified MtrA. **B:** Mutating either of the two most conserved nucleotides within this sequence (T_1_ or C_4_) significantly reduced MtrA binding to DNA.

### D53 is a key residue for MtrA DNA binding

Upon activation by an environmental signal, MtrB is predicted to autophosphorylate using ATP and then phosphorylate MtrA on residue D53 (39). In some (but not all) RRs changing this aspartate to glutamate (E) can mimic phosphorylation and activate the protein whereas changing it to alanine (A) inactivates typical RRs (40). To investigate the binding of MtrA to DNA and its dependence on phosphorylation, wild-type, D53A and D53E MtrA proteins were over-produced and purified from *E. coli*. Using the *mtrA* promoter probe, the DNA binding affinities of freshly purified wild-type, D53A and D53E MtrA proteins were measured *in vitro* using ReDCaT SPR. This revealed that wild-type MtrA has the strongest binding but D53E MtrA retains some DNA binding activity to the *mtrA* promoter probe at higher concentrations. The D53A MtrA variant did not bind to this DNA probe (Figure 3).

**Figure 3.**
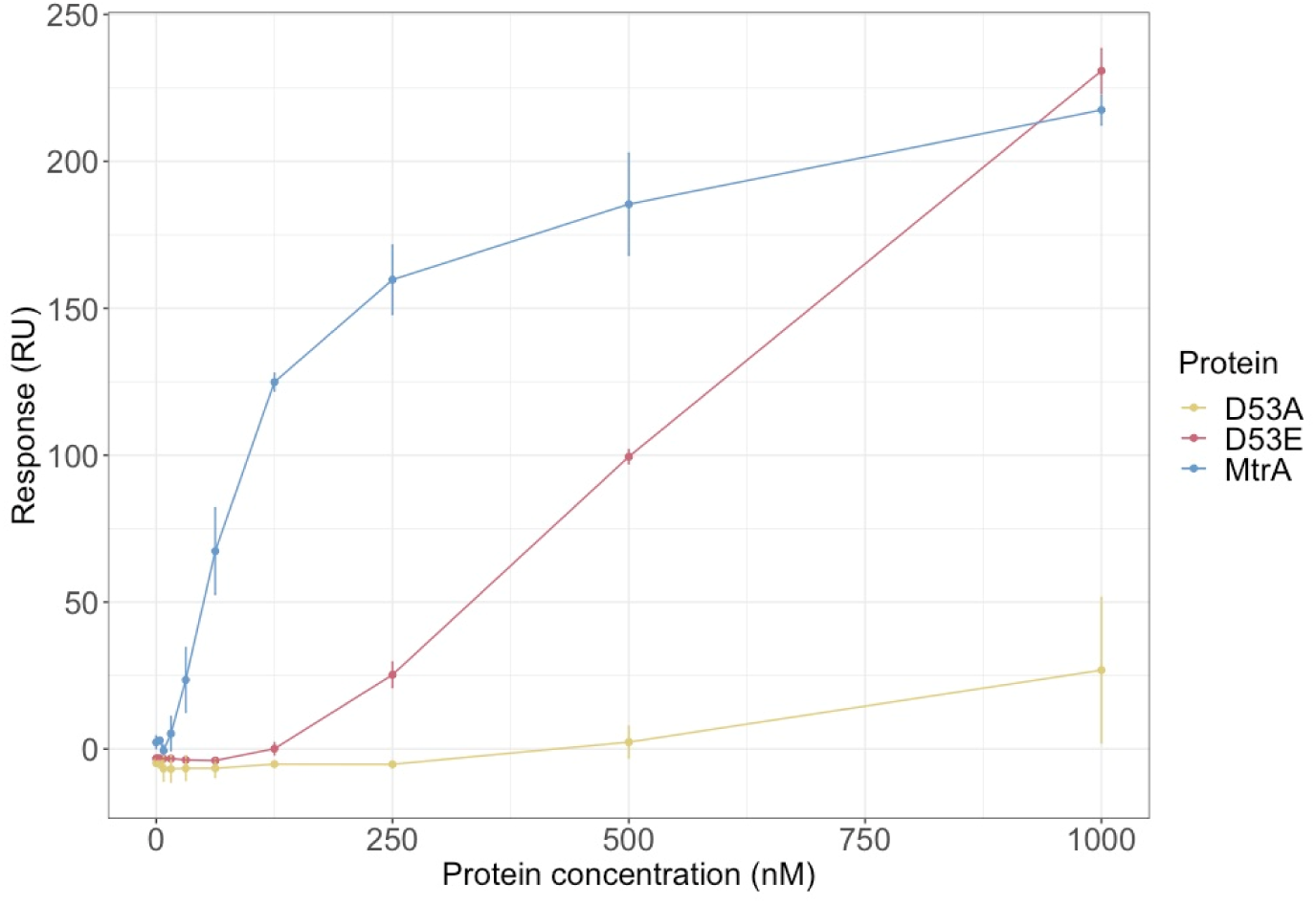
DNA binding by MtrA requires D53. Altering D53 to glutamate (D53E) reduces the ability of MtrA to bind to the *mtrA* promoter while changing it to alanine (D53A) abolishes DNA binding activity.

A recent study using *S. venezuelae* ATCC-10712 reported that a D53E change prevents MtrA from binding to DNA in electrophoretic mobility shift assays (EMSAs) and that complementing an Δ*mtrA* mutant *in trans* with alleles encoding Flag-tagged D53A, D53E and D53N variants failed to pull down known MtrA target promoters in a ChIP-PCR experiment (39). The divergence in our results may be because they used different promoters or because ReDCaT SPR is a more sensitive way of measuring *in vitro* DNA binding activity than EMSAs. We conclude from our data that D53 is the site of phosphorylation in *S. venezuelae* NRRL B-65442 and that the D53E MtrA variant retains some DNA binding activity, at least *in vitro*.

### MtrA directly controls the expression of key developmental regulatory genes

Next we used tandem mass tag (TMT) proteomics to examine the effects of *mtrA* deletion on the expression of developmental genes when the wild-type and Δ*mtrA* strains were grown on R2YE and MYM agar (Figure 4; Table 5, Table S2). Despite having MtrA binding sites, the expression of the *adpA*, *dnaA*, *ssgB* and *filP* genes was not significantly affected by loss of MtrA on MYM or R2YE agar. However, the expression of *bldM* and *whiI* were significantly affected on R2YE but not MYM agar, with BldM levels 4-fold lower and WhiI levels 7-fold lower in the Δ*mtrA* mutant grown on R2YE agar (Table 5). This suggests that MtrA directly activates the expression of both *bldM* and *whiI* on R2YE agar and that loss of MtrA leads to a reduction of BldM and WhiI that could be responsible for the bald phenotype of the Δ*mtrA* strain (Figure 1).

**Figure 4.**
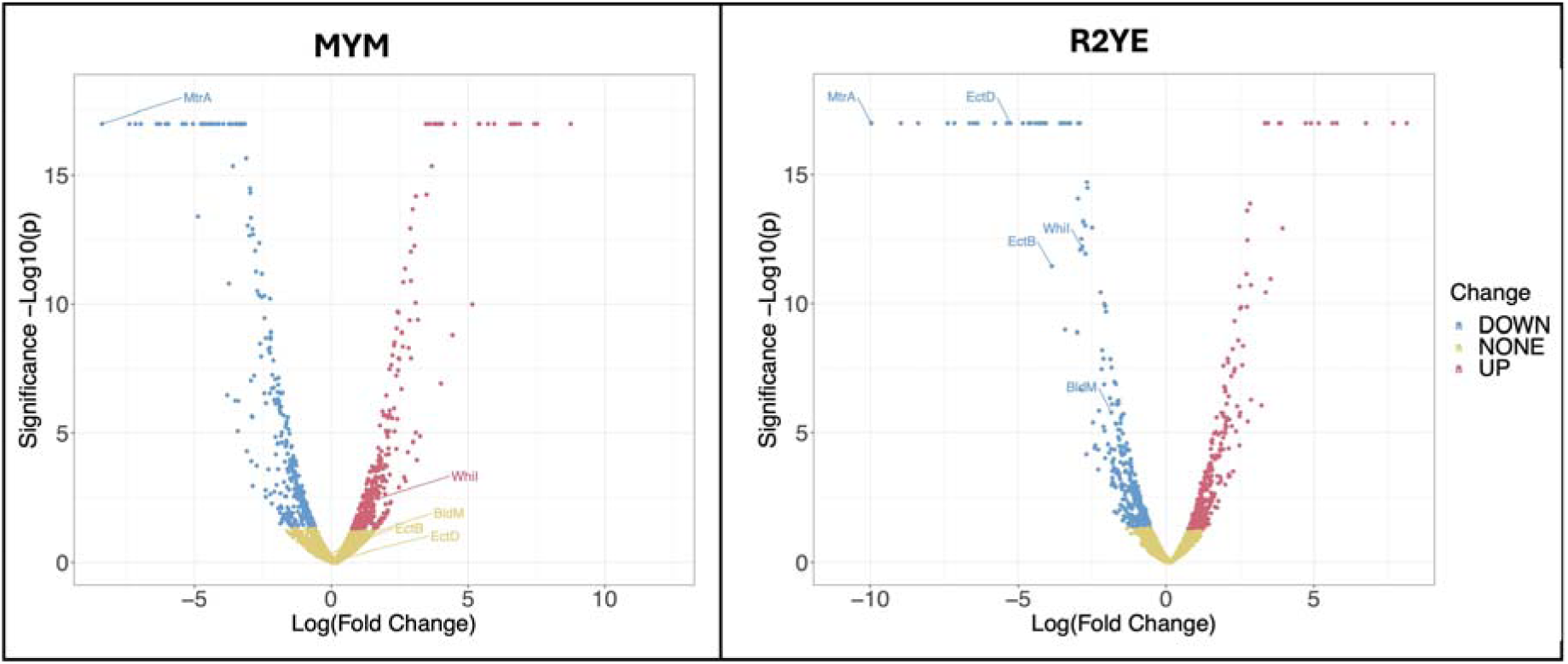
Loss of MtrA results in significantly lower levels of BldM and WhiI on R2YE agar. TMT proteomics of *S. venezualae* shows that, when grown on R2YE (without glucose) the Δ*mtrA* mutant has significantly lower levels of the developmental regulators BldM and WhiI and the ectoine biosynthetic enzymes EctB and EctD (highlighted with arrows on the vocano plot).

BldM and WhiI are key regulators of development and control the expression of genes required for aerial hyphae formation and sporulation (Table 5) (30). As such, a *bldM* mutant is bald because it does not produce aerial hyphae and a *whiI* mutant is white because it does not produce spores and therefore lacks the green WhiE spore-pigment. They are both atypical response regulators because they do not require phosphorylation to bind to DNA. Homodimeric BldM regulates class I BldM target genes which are required for the production of aerial hyphae while heterodimeric BldM-WhiI regulates class II BldM target genes that are required for sporulation (30). BldM activates the expression of *ssgR* which encodes a transcription factor that in turn activates the expression of *ssgA* which is required to correctly position Z rings prior to cell division (sporulation) (30). Consistent with this, SsgR levels are 4-fold lower in the MtrA mutant growing on R2YE agar and SsgA levels are 2-fold lower but their levels are not significantly affected on MYM agar (Table 5). Homodimeric BldM also activates *whiB* expression which is 2-fold lower in the Δ*mtrA* mutant grown on R2YE but not MYM agar (Table 5). BldM-WhiI targets include the *whiE* operon (30) whose gene products make the green spore-pigment and are down-regulated up to 3-fold in the Δ*mtrA* mutant grown on R2YE but not MYM agar (Table 5). Thus, the combined TMT proteomics, ChIP-seq and ReDCaT SPR data are consistent with MtrA directly activating the production of BldM and WhiI and the expression of their downstream class I and class II target genes on R2YE agar. We hypothesise that loss of MtrA reduces the levels of BldM and WhiI under these growth conditions and result in the observed Δ*mtrA* bald phenotype.

### Evidence that MtrAB senses osmotic stress

The conditional bald phenotype of Δ*mtrA* mutants in *Streptomyces* species suggests that MtrA is only required to activate entry into sporulation under certain growth conditions. The sensor kinase MtrB has been reported to sense osmotic stress in *C. glutamicum* and activate MtrA which in turn activates the expression of *betP* and *proP* which encode transporters for the compatible solutes betaine and proline (41,42). The reported *Streptomyces* Δ*mtrA* mutants are all bald on R2YE agar and it is notable that R2YE contains 10.2% sucrose which likely induces osmotic stress. To test whether sucrose is responsible for the bald phenotype we grew the wild-type, Δ*mtrA* and complemented Δ*mtrA* strains on R2YE and MYM agar with and without 10.2% sucrose. The results show that the *S. venezuelae* NRRL B65442 Δ*mtrA* mutant is only bald on R2YE agar that contains sucrose and that adding 10.2% sucrose to MYM agar delays sporulation in the Δ*mtrA* mutant (Figure 5).

**Figure 5.**
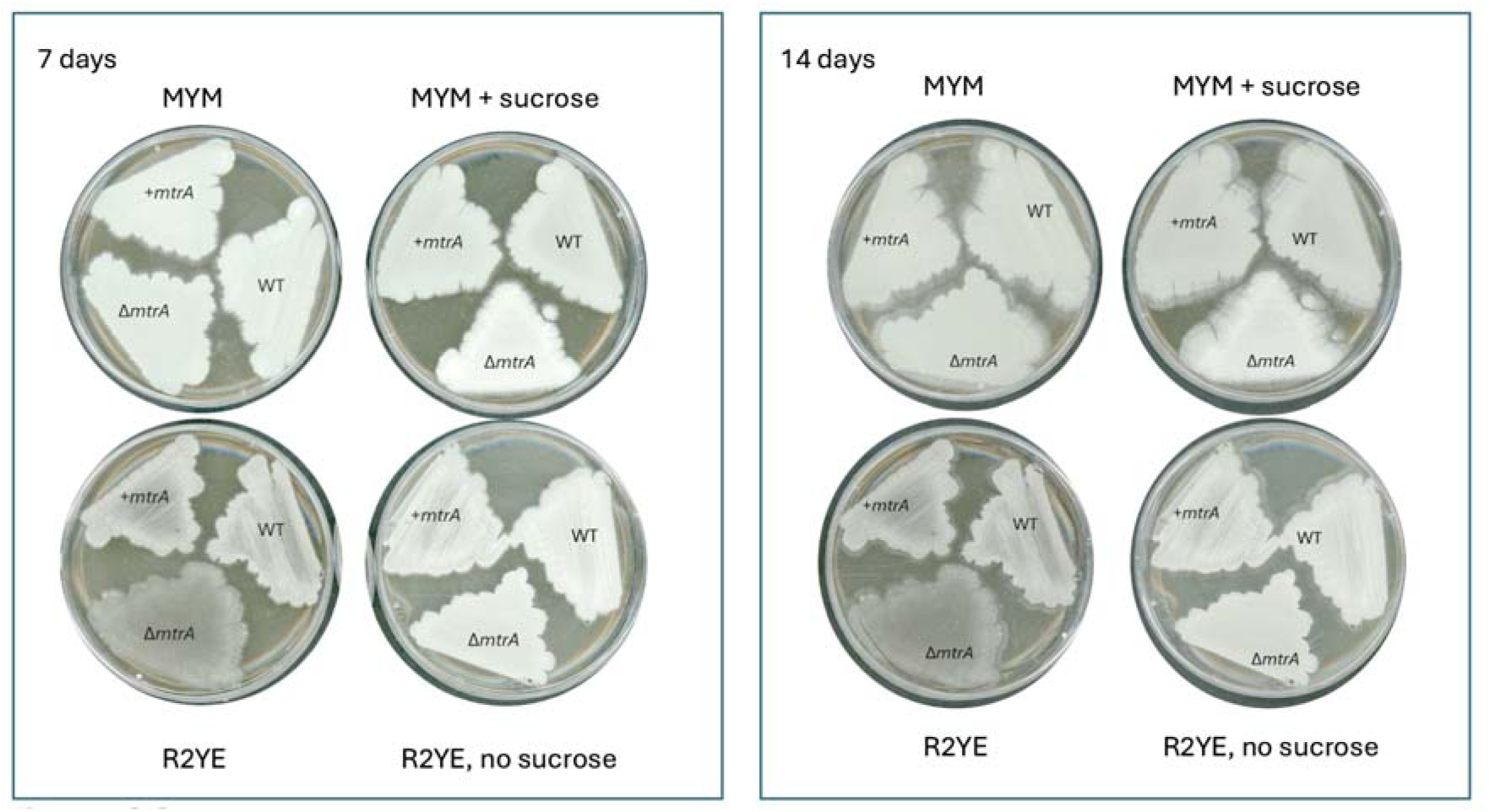
Sucrose induces the Δ*mtrA* bald phenotype. *S. venezuelae* wild-type, Δ*mtrA* and Δ*mtrA* + *mtrA* (complemented) strains grown on MYM and R2YE agar with and without 10.2% sucrose for 7 and 14 days.

These results indicate that MtrAB controls the expression of its target genes in response to the high level of sucrose in the growth medium, and we hyopothesise that this sucrose is inducing osmotic stress. Consistent with this, the *ectABCD* operon promoter is highly enriched in the MtrA ChIP-seq dataset throughout the developmental time course (Figure 6A). This operon encodes the biosynthetic pathway for the compatible solute and osmoprotectant ectoine. EctA and EctC were not detected in the TMT experiment, but the data show that EctB and EctD were downregulated 14- and 38-fold, respectively, in the Δ*mtrA* mutant relative to wild-type on R2YE agar (Figure 6B, Table S3). Notably, the levels of these proteins were not significantly affected in Δ*mtrA* versus wild-type grown on MYM agar (0.797 and 1.246, respectively) which further supports our hypothesis that MtrAB senses and responds to osmotic stress induced by 10.2% sucrose in R2YE agar. Extraction and LCMS analysis of the *S. venezuelae* wild-type and mutant strains grown on MYM or R2YE agar with and without sucrose show a significant increase in ectoine production in the wild-type strain in the presence of 10.2% sucrose and this is abolished in the Δ*mtrA* mutant (Figure 6C). Taken together the data presented in Figure 6 indicate that MtrA directly regulates ectoine biosynthesis in response to high sucrose concentrations. Given that ectoine is produced to protect *Streptomyces* bacteria against heat and osmotic stress, we propose that MtrAB is a conserved osmotic stress sensing two-component system in the genus *Streptomyces*.

**Figure 6.**
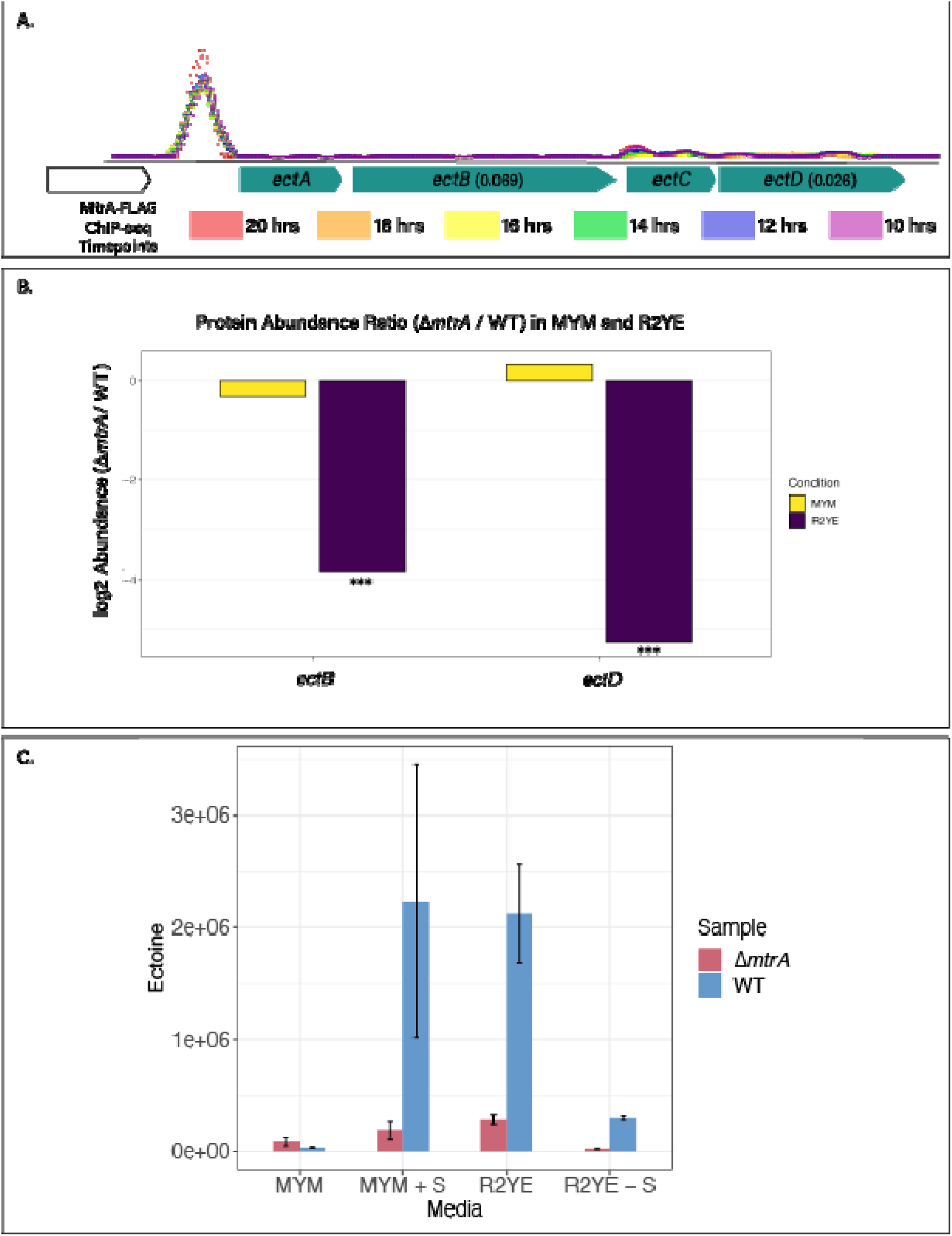
MtrA directly activates ectoine biosynthesis. (A). ChIP-seq data showing that MtrA binds to the *ectABCD* operon promoter throughout a developmental time course. (B). TMT proteomics data showing that the operon products EctB and EctD are 14- and 38-fold lower in the Δ*mtrA* mutant relative to wild-type (WT) suggesting MtrA directly activates the expression of the *ectABCD* operon. EctA and EctC were not detected in this experiment. (C) LCMS data showing the levels of ectoine produced in *S. venezuelae* WT and Δ*mtrA* strains grown on MYM and R2YE agar with and without 10.2% sucrose (+S or –S).

### Over-expression of *bldM* and *whiI* restores normal development in the Δ*mtrA* mutant

To test whether the reduced levels of BldM and WhiI are responsible for the conditional bald phenotype of the Δ*mtrA* mutant under osmotic stress conditions, we over-expressed the *bldM* and *whiI* genes in the Δ*mtrA* mutant using the constitutive high level *ermE\** promoter. These strains were then grown on MYM with and without 10.2% sucrose. The results show that in the presence of sucrose, the *mtrA* mutant is unable to sporulate, but this is complemented by the overexpression of *bldM* and *whiI* (Figure 7). This supports our hypothesis that MtrAB activates the expression of *bldM* and *whiI* under osmotic stress conditions (i.e., in the presence of 10.2% sucrose) to trigger sporulation and that deletion of *mtrA* leads to a reduction in BldM and WhiI levels under these growth conditions and prevents sporulation, resulting in a conditional bald phenotype.

**Figure 7.**
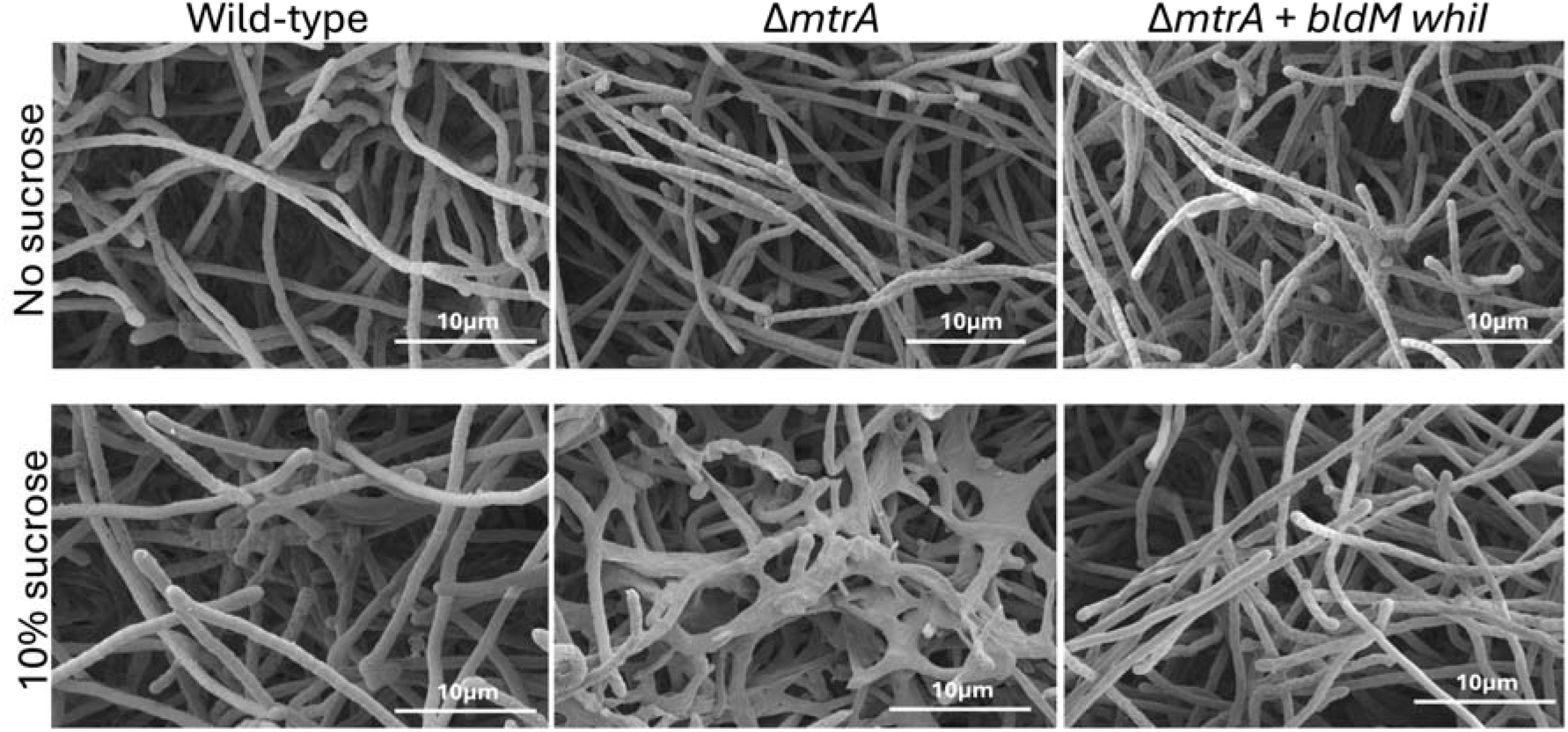
Over-expression of *bldM* and *whiI* in the Δ*mtrA* mutant. Scanning electron micrographs were taken after 10 days growth on MYM agar with or without 10.2% sucrose, as indicated. The images show that even after this extended incubation, *S. venezuelae* Δ*mtrA* is unable to develop normal spore chains in the presence of 10% sucrose. However, artificial overexpression of *bldM* and *whiI* in a synthetic operon under the control of the constitutive high level *ermE\** promoter restores sporulation to the Δ*mtrA* mutant under osmotic stress conditions.

This work further illustrates the complexity of *Streptomyces* life cycle regulation, where sporulation must be controlled temporally and in response to both biotic and abiotic environmental stresss. BldM and WhiI act together to initiate sporulation (30) and both are under the direct or indirect control of BldD, the master repressor of sporulation (43). BldD directly represses the expression of *bldM* and *whiG*, the latter encoding the RNA polymerase sigma factor σ^WhiG^, which directs transcription of *whiI* (10). σ^WhiG^ activity is controlled post-translationally by the anti-σ factor RsiG (44) and both BldD and RsiG are controlled post-translationally by binding of the secondary messenger cyclic-di-GMP (10,45). Thus, a reduction in intracellular c-di-GMP levels (in response to unknown signals) relieves BldD repression of *bldM* and *whiG* and removes RsiG inhibition of σ^WhiG^, leading to σ^WhiG^-directed transcription of *whiI* (46). It is not clear how MtrA bypasses BldD and RsiG repression to simultaneously activate the expression of both *bldM* and *whiI* or if MtrA-dependent activation of *whiI* requires σ^WhiG^. These are all questions for future studies but the levels of BldD, σ^WhiG^ and RsiG were not significantly affected by the loss of MtrA on either MYM or R2YE agar in this work (Table S2). Our ReDCaT SPR experiments show that purified MtrA can bind to five different sites at the *bldM* promoter, three of which are upstream of the transcript start site (TSS), with two downstream (Table 1). Only one is positioned (-42 relative to the TSS) such that bound MtrA could contact bound RNA polymerase to activate *bldM* expression. MtrA binds to only single site at the *whiI* promoter, and this is positioned at -55 relative to the TSS, which is consistent with MtrA-dependent activation via contact with bound RNA polymerase. It will be important to understand which of the sites at the *bldM* promoter are bound by MtrA *in vivo* under osmotic stress conditions and if or how MtrA interacts with BldD and σ^WhiG^. It has been reported that the *bldD* promoter is directly repressed by MtrA in *S. avermitilis* and *S. coelicolor* (22,29), but *bldD* was not identified as an MtrA target in ChIP-seq experiments in *S. venezuelae* NRRL B-65442 (1), and the *bldD* promoter was not bound by purified MtrA in the ReDCaT SPR experiments.

## Concluding Remarks

Here we have shown that the conditional bald phenotype of the *S. venezuelae* Δ*mtrA* mutant is caused by the addition of 10.2% sucrose to the growth medium. Under these conditions the Δ*mtrA* mutant fails to form aerial hyphae and spores. Given that high levels of sucrose are known to induce hyperosmotic stress in *Streptomyces* bacteria (47,48) we propose that MtrAB is an osmotic stress sensing two component system. In our model, the sensor kinase MtrB senses osmotic stress, autophosphorylates and then phosphorylates its cognate response regulator MtrA at residue D53 to activate its DNA binding activity. MtrA then responds to the osmotic stress by activating the production of the compatible solute ectoine and triggering entry into sporulation via activation of BldM and WhiI, thus increasing the chances of survival (30). *Streptomyces* species live in constantly fluctuating soil environments where osmotic stress is likely to present a frequent danger. Loss of *mtrA* has been shown to result in bald phenotypes on R2YE agar in distantly related *Streptomyces* species and we propose that the osmotic stress sensing function of MtrAB is highly conserved in this genus. Furthermore, MtrAB has been shown to sense and respond to osmotic stress in *C. glutamicum* and *Dietzia* spp, suggesting MtrAB function is even more widely conserved in the phylum Actinomycetota (21,42,49).

## Supporting information

Proteomics data

## Author statements

### Author contributions

*Conceptualization – ADMB, RD, NFS, BW, MIH*

*Data Curation – ADMB, GS, NFS, RD,*

*Formal Analysis – ADMB, GS NFS, RD*

*Funding Acquisition – BW, MIH*

*Investigation –ADMB, GS, KN, MR, NFS, NH, RS, RD,*

*Methodology – LB, NAH, NFS, RD*

*Project Administration – ADMB, BW, NFS, RD, MIH*

*Resources – BW, GS, MIH, MR, RS*

*Supervision –BW, MIH*

*Validation – ADMB, NFS, RD*

*Visualization – NFS, RD, ADMB*

*Writing – Original Draft – MIH*

*Writing – Review & Editing – ADMB, BW, KN, LB, NFS, NAH, RD, GS, MIH*

### Conflicts of interest

The authors declare that there are no conflicts of interest.

### Funding information

Nicolle Som: BBSRC doctoral training programme grant BB/M011216/1 and responsive mode grant BB/P005292/1. Rebecca Devine: BBSRC doctoral training programme grant BB/M011216/1 and responsive mode grant BB/W000628/1. Ainsley Beaton: responsive mode grant BB/W000628/1. Gerhard Saalbach and Neil Holmes: BBSRC-funded Institute Strategic Program Harnessing Biosynthesis for Sustainable Food and Health (HBio) grant BB/X01097X/1 to the John Innes Centre. The John Innes Centre Bioimaging Platform is supported by the UKRI Biotechnology and Biological Sciences Research Council (grant BB/CCG2240/1).

### Ethical approval

Not applicable

## Acknowledgements

We thank our colleagues Mark Buttner and Matt Bush for critical reading of the manuscript and useful discussion and suggestions. We also thank past and present members of the Hutchings and Wilkinson groups at the John Innes Centre (JIC) and JIC support services for growth media, glassware, plasticware and waste disposal.

## Notes

### Competing Interest Statement

The authors have declared no competing interest.

## References

1. Som NF, Heine D, Holmes NA, Munnoch JT, Chandra G, Seipke RF, et al. The Conserved Actinobacterial Two-Component System MtrAB Coordinates Chloramphenicol Production with Sporulation in Streptomyces venezuelae NRRL B-65442. Front Microbiol. 2017 June 28;8:1145.

2. Munnoch JT, Martinez MTP, Svistunenko DA, Crack JC, Le Brun NE, Hutchings MI. Characterization of a putative NsrR homologue in Streptomyces venezuelae reveals a new member of the Rrf2 superfamily. Sci Rep. 2016 Sept 8;6(1):31597.

3. Feeney MA, Newitt JT, Addington E, Algora-Gallardo L, Allan C, Balis L, et al. ActinoBase: tools and protocols for researchers working on Streptomyces and other filamentous actinobacteria. Microb Genomics. 2022 Jan 1;8(7):mgen000824.

4. Hutchings MI, Truman AW, Wilkinson B. Antibiotics: past, present and future. Curr Opin Microbiol. 2019 Oct;51:72–80.

5. Schlimpert S, Elliot MA. The Best of Both Worlds—Streptomyces coelicolor and Streptomyces venezuelae as Model Species for Studying Antibiotic Production and Bacterial Multicellular Development. J Bacteriol. 2023 June 22;e00153–23.

6. Newitt J, Prudence S, Hutchings M, Worsley S. Biocontrol of Cereal Crop Diseases Using Streptomycetes. Pathogens. 2019 June 13;8(2):78.

7. Worsley SF, Newitt J, Rassbach J, Batey SFD, Holmes NA, Murrell JC, et al. *Streptomyces* Endophytes Promote Host Health and Enhance Growth across Plant Species. Appl Environ Microbiol. 2020 Aug 3;86(16):e01053–20.

8. Viaene T, Langendries S, Beirinckx S, Maes M, Goormachtig S. *Streptomyces* as a plant’s best friend? FEMS Microbiol Ecol. 2016 Aug;92(8):fiw119.

9. Dow L, Gallart M, Ramarajan M, Law SR, Thatcher LF. Streptomyces and their specialised metabolites for phytopathogen control – comparative in vitro and in planta metabolic approaches. Front Plant Sci. 2023 June 14;14:1151912.

10. Bush MJ, Tschowri N, Schlimpert S, Flärdh K, Buttner MJ. c-di-GMP signalling and the regulation of developmental transitions in streptomycetes. Nat Rev Microbiol. 2015 Dec;13(12):749–60.

11. Hutchings MI, Wilkinson B. Insects and their antibiotic-producing bacteria. Microbiota Host. 2023 Oct 6;1(1):e230008.

12. Seipke RF, Kaltenpoth M, Hutchings MI. *Streptomyces* as symbionts: an emerging and widespread theme? FEMS Microbiol Rev. 2012 July;36(4):862–76.

13. Baltz RH. Gifted microbes for genome mining and natural product discovery. J Ind Microbiol Biotechnol. 2017 May 1;44(4–5):573–88.

14. McLean TC, Lo R, Tschowri N, Hoskisson PA, Al Bassam MM, Hutchings MI, et al. Sensing and responding to diverse extracellular signals: an updated analysis of the sensor kinases and response regulators of Streptomyces species. Microbiology. 2019 Sept 1;165(9):929–52.

15. Hoskisson PA, Hutchings MI. MtrAB–LpqB: a conserved three-component system in actinobacteria? Trends Microbiol. 2006 Oct;14(10):444–9.

16. Nguyen HT, Wolff KA, Cartabuke RH, Ogwang S, Nguyen L. A lipoprotein modulates activity of the MtrAB two-component system to provide intrinsic multidrug resistance, cytokinetic control and cell wall homeostasis in *Mycobacterium*. Mol Microbiol. 2010 Apr;76(2):348–64.

17. Zahrt TC, Deretic V. An Essential Two-Component Signal Transduction System in *Mycobacterium tuberculosis*. J Bacteriol. 2000 July;182(13):3832–8.

18. Chatterjee A, Sharma AK, Mahatha AC, Banerjee SK, Kumar M, Saha S, et al. Global mapping of MtrA-binding sites links MtrA to regulation of its targets in Mycobacterium tuberculosis. Microbiology. 2018 Jan 1;164(1):99–110.

19. Plocinska R, Purushotham G, Sarva K, Vadrevu IS, Pandeeti EVP, Arora N, et al. Septal Localization of the Mycobacterium tuberculosis MtrB Sensor Kinase Promotes MtrA Regulon Expression. J Biol Chem. 2012 July;287(28):23887–99.

20. Purushotham G, Sarva KB, Blaszczyk E, Rajagopalan M, Madiraju MV. Mycobacterium tuberculosis oriC sequestration by MtrA response regulator: MtrA and M. tuberculosis cell cycle. Mol Microbiol. 2015 Oct;98(3):586–604.

21. Krämer R. Osmosensing and osmosignaling in Corynebacterium glutamicum. Amino Acids. 2009 Sept;37(3):487–97.

22. Zhang P, Wu L, Zhu Y, Liu M, Wang Y, Cao G, et al. Deletion of MtrA Inhibits Cellular Development of Streptomyces coelicolor and Alters Expression of Developmental Regulatory Genes. Front Microbiol. 2017 Oct 16;8:2013.

23. Zhu Y, Zhang P, Zhang J, Xu W, Wang X, Wu L, et al. The developmental regulator MtrA binds GlnR boxes and represses nitrogen metabolism genes in *Streptomyces coelicolor*. Mol Microbiol. 2019 July;112(1):29–46.

24. Zhu Y, Wang J, Su W, Lu T, Li A, Pang X. Effects of dual deletion of *glnR* and *mtrA* on expression of nitrogen metabolism genes in *Streptomyces venezuelae*. Microb Biotechnol. 2022 June;15(6):1795–810.

25. Parra J. Antibiotics from rare actinomycetes, beyond the genus Streptomyces. Curr Opin Microbiol. 2023;

26. Pan Q, Tong Y, Han YJ, Ye BC. Two amino acids missing of MtrA resulted in increased erythromycin level and altered phenotypes in Saccharopolyspora erythraea. Appl Microbiol Biotechnol. 2019 June;103(11):4539–48.

27. Som NF, Heine D, Holmes N, Knowles F, Chandra G, Seipke RF, et al. The MtrAB two-component system controls antibiotic production in Streptomyces coelicolor A3(2). Microbiology. 2017 Oct 1;163(10):1415–9.

28. Zhu Y, Zhang P, Zhang J, Wang J, Lu Y, Pang X. Impact on Multiple Antibiotic Pathways Reveals MtrA as a Master Regulator of Antibiotic Production in Streptomyces spp. and Potentially in Other Actinobacteria. Appl Environ Microbiol. 2020 Oct;86(20):e01201–20.

29. Tian J, Li Y, Zhang C, Su J, Lu W. Characterization of a pleiotropic regulator MtrA in Streptomyces avermitilis controlling avermectin production and morphological differentiation. Microb Cell Factories. 2024 Apr 8;23(1):103.

30. Al-Bassam MM, Bibb MJ, Bush MJ, Chandra G, Buttner MJ. Response Regulator Heterodimer Formation Controls a Key Stage in Streptomyces Development. PLoS Genet. 2014 Aug 7;10(8):e1004554.

31. Bursy J, Kuhlmann AU, Pittelkow M, Hartmann H, Jebbar M, Pierik AJ, et al. Synthesis and Uptake of the Compatible Solutes Ectoine and 5-Hydroxyectoine by *Streptomyces coelicolor* A3(2) in Response to Salt and Heat Stresses. Appl Environ Microbiol. 2008 Dec;74(23):7286–96.

32. Cobb RE, Wang Y, Zhao H. High-Efficiency Multiplex Genome Editing of *Streptomyces* Species Using an Engineered CRISPR/Cas System. ACS Synth Biol. 2015 June 19;4(6):723–8.

33. Gregory MA, Till R, Smith MCM. Integration Site for Streptomyces Phage φBT1 and Development of Site-Specific Integrating Vectors. J Bacteriol. 2003 Sept;185(17):5320–3.

34. Schlimpert S, Wasserstrom S, Chandra G, Bibb MJ, Findlay KC, Flärdh K, et al. Two dynamin-like proteins stabilize FtsZ rings during Streptomyces sporulation. Proc Natl Acad Sci. 2017 July 25;114(30):E6176–83.

35. Hong HJ, Hutchings MI, Hill LM, Buttner MJ. The Role of the Novel Fem Protein VanK in Vancomycin Resistance in Streptomyces coelicolor. J Biol Chem. 2005 Apr;280(13):13055–61.

36. Brenes A, Hukelmann J, Bensaddek D, Lamond AI. Multibatch TMT Reveals False Positives, Batch Effects and Missing Values. Mol Cell Proteomics. 2019 Oct;18:1967–80.

37. Eng JK, Jahan TA, Hoopmann MR. Comet: An open-source MS / MS sequence database search tool. Proteomics. 2013 Jan;13(1):22–4.

38. Stevenson CEM, Lawson DM. Analysis of Protein–DNA Interactions Using Surface Plasmon Resonance and a ReDCaT Chip. In: Daviter T, Johnson CM, McLaughlin SH, Williams MA, editors. Protein-Ligand Interactions: Methods and Applications. New York, NY: Springer US; 2021. p. 369–79.

39. Lu T, Zhu Y, Ni X, Zhang X, Liu Y, Cui X, et al. Mutation of MtrA at the Predicted Phosphorylation Site Abrogates Its Role as a Global Regulator in Streptomyces venezuelae. Microbiol Spectr. 2022 Apr 27;10(2):e02131–21.

40. Devine R, Noble K, Stevenson C, De Oliveira Martins C, Saalbach G, McDonald HP, et al. Redox control of antibiotic biosynthesis. mBio. 2025 Aug;16(9):e0136925

41. Möker N, Reihlen P, Krämer R, Morbach S. Osmosensing Properties of the Histidine Protein Kinase MtrB from. J Biol Chem. 2007 Sept;282(38):27666–77.

42. Möker N, Brocker M, Schaffer S, Krämer R, Morbach S, Bott M. Deletion of the genes encoding the MtrA–MtrB two-component system of *Corynebacterium glutamicum* has a strong influence on cell morphology, antibiotics susceptibility and expression of genes involved in osmoprotection. Mol Microbiol. 2004 Oct;54(2):420–38.

43. Den Hengst CD, Tran NT, Bibb MJ, Chandra G, Leskiw BK, Buttner MJ. Genes essential for morphological development and antibiotic production in Streptomyces coelicolor are targets of BldD during vegetative growth: The Streptomyces BldD regulon. Mol Microbiol. 2010 Oct;78(2):361–79.

44. Gallagher KA, Schumacher MA, Bush MJ, Bibb MJ, Chandra G, Holmes NA, et al. c-di-GMP Arms an Anti-σ to Control Progression of Multicellular Differentiation in Streptomyces. Mol Cell. 2020 Feb;77(3):586–599.e6.

45. Tschowri N, Schumacher MA, Schlimpert S, Chinnam N babu, Findlay KC, Brennan RG, et al. Tetrameric c-di-GMP Mediates Effective Transcription Factor Dimerization to Control Streptomyces Development. Cell. 2014 Aug;158(5):1136–47.

46. Gallagher KA, Tschowri N, Brennan RG, Schumacher MA, Buttner MJ. How c-di-GMP controls progression through the Streptomyces life cycle. Curr Opin Microbiol. 2024 Aug;80:102516.

47. Alonso-Fernández S, Gutiérrez-Del-Río I, Lombó F, Fernández-Del-Campo-García MT, Herrero-Hernández E, García-Gómez D, et al. Peptidoglycan-reshuffling proteins SCO0954, SCO1758, SCO4439, and SCO4440 modulate the formation of wall-deficient cells in Streptomyces coelicolor under hyperosmotic sucrose stress. Sci Rep. 2025 Sept 1;15(1):32112.

48. Bhowmick S, Shenouda ML, Tschowri N. Osmotic stress responses and the biology of the second messenger c-di-AMP in Streptomyces. MicroLife. 2023 Jan 4;4:uqad020.

49. Qin X, Zhang K, Nie Y, Wu XL. The Roles of the Two-Component System, MtrAB, in Response to Diverse Cell Envelope Stresses in *Dietzia* sp. DQ12-45-1b. Zhou NY, editor. Appl Environ Microbiol. 2022 Oct 26;88(20):e01337–22.

