## Supplementary data for "MtrAB activates ectoine production and triggers sporulation in response to osmotic stress in *Streptomyces venezuelae*"

**Table S1. Primers used in this study**

| **Primer name** | **Description** | **Sequence** |
| --- | --- | --- |
| NS-145 | *mtrA* forward BamHI (pETduet) | GGATCCTAAAGGCCGCGTTCTGGTCGTC |
| NS-146 | *mtrA* reverse HindIII (pETduet) | AAGCTT TCA GCTCGGTCCCGCCTTGTACC |
| NS301 | *mtrA* forward HindIII (pSS170) | ccatcagcaaaaggggatgataagtttatcCCGTGGGCCGGTCCC |
| NS301 | *mtrA* reverse HindIII (pSS170) | CGAATCGATGATCATATGAGAGAATCTtcagctcggtcccgcctt |
| NS304 | *mtrA* left arm in pCRISP | gaacgctcggttgccgccgggcgttttttaGACCGGGCGGCCGAG |
| NS305 | *mtrA* left arm in pCRISP | ccacgatctccgggcgctccgggtcATCGACATCATCCCATTTCCGTATC |
| NS306 | *mtrA* right arm in pCRISP | AGACTGATACGGAAATGGGATGATGTCGATgacccggagcgcccg |
| NS307 | *mtrA* right arm in pCRISP | ttttacggttcctggccgcgccggggccgCTCGCCGGTGACCTTCATCC |
| NS308 | *mtrA* spacer 1 For | ACGCCGACCTGATGCTGCCCGGAA |
| NS309 | *mtrA* spacer 1 Rev | AAACTTCCGGGCAGCATCAGGTCG |
| NS365 | *pETduet D53 For GA* | ccacagccaggatccTAAAGGCCGCGTTCTGGTCGTCGACGACGACACCG |
| NS366 | *pETduet D53 Rev GA* | TTAAGCATTATGCGGCCGCAAGCTTtcagctcggtcccgcctt |
| RD001 | *bldM* for NdeI | gtctagaacaggaggccccatatgATGACATCCGTTCTCGTC |
| RD002 | *bldM* rev plus RBS | TTACCTCCGATGTTGAGTGCGATTCAGCGAACCAG |
| RD003 | *whiI* For plus RBS | CTCAACATCGGAGGTAAGCCATGTCCGTTCTCCTCGAGC |
| RD004 | *whiI* Rev HindIII | ctcatgagaacctaggatccaagcttGTCCGTCAGTGGATGATC |

**Table S2. Oligos used for ReDCaT SPR.** Upper case represents promoter sequence, lower case is the ReDCaT linker for annealing to the chip.

| **Primer name** | **Sequence** |
| --- | --- |
| ssgBFor_1 | GGTCCCAACTTACCCGACGGCGGGCAGTCGCCGGAATGGC |
| ssgBRev_1 | GCCATTCCGGCGACTGCCCGCCGTCGGGTAAGTTGGGACCcctaccctacgtcctcctgc |
| ssgBFor_2 | AGTCGCCGGAATGGCGGTATTACCGCAGGTCACACGCCCC |
| ssgBRev_2 | GGGGCGTGTGACCTGCGGTAATACCGCCATTCCGGCGACTcctaccctacgtcctcctgc |
| ssgBFor_3 | CAGGTCACACGCCCCCATACGAGCTACCTGAACGATCACA |
| ssgBRev_3 | TGTGATCGTTCAGGTAGCTCGTATGGGGGCGTGTGACCTGcctaccctacgtcctcctgc |
| ssgBFor_4 | ACCTGAACGATCACAAAACCGACAGAGGGGTTTACAACGG |
| ssgBRev_4 | CCGTTGTAAACCCCTCTGTCGGTTTTGTGATCGTTCAGGTcctaccctacgtcctcctgc |
| ssgBFor_5 | AGGGGTTTACAACGGCACTGGAGGTGGCATGTCGATTTCG |
| ssgBRev_5 | CGAAATCGACATGCCACCTCCAGTGCCGTTGTAAACCCCTcctaccctacgtcctcctgc |
| ssgBFor_6 | GGCATGTCGATTTCGCCGACGTGCGAATCCCCGAGCGCAC |
| ssgBRev_6 | GTGCGCTCGGGGATTCGCACGTCGGCGAAATCGACATGCCcctaccctacgtcctcctgc |
| ssgBFor_7 | AATCCCCGAGCGCACACTGAGCGAAAGGCCCTGGCGCTTA |
| ssgBRev_7 | TAAGCGCCAGGGCCTTTCGCTCAGTGTGCGCTCGGGGATTcctaccctacgtcctcctgc |
| ssgBFor_8 | AGGCCCTGGCGCTTATGAACACCAC |
| ssgBRev_8 | GTGGTGTTCATAAGCGCCAGGGCCTcctaccctacgtcctcctgc |
| adpAFor_1 | GGGCCCGGCCCACGGGGCGGACCCCGGGGGAGAAACCGGT |
| adpaARev_1 | ACCGGTTTCTCCCCCGGGGTCCGCCCCGTGGGCCGGGCCCcctaccctacgtcctcctgc |
| adpAFor_1 | GGGCCCGGCCCACGGGGCGGACCCCGGGGGAGAAACCGGT |
| adpARev_1 | ACCGGTTTCTCCCCCGGGGTCCGCCCCGTGGGCCGGGCCCcctaccctacgtcctcctgc |
| adpAFor_2 | GGGGGAGAAACCGGTATCAATCGGGGCGCCGACGGGGCCC |
| adpARev_2 | GGGCCCCGTCGGCGCCCCGATTGATACCGGTTTCTCCCCCcctaccctacgtcctcctgc |
| adpAFor_3 | GCGCCGACGGGGCCCGCTCCTCGTGGAGTTGACAGGTTCG |
| adpARev_3 | CGAACCTGTCAACTCCACGAGGAGCGGGCCCCGTCGGCGCcctaccctacgtcctcctgc |
| adpAFor_4 | GAGTTGACAGGTTCGTACACCCGAACCCATATGTTTTCAG |
| adpARev_4 | CTGAAAACATATGGGTTCGGGTGTACGAACCTGTCAACTCcctaccctacgtcctcctgc |
| adpAFor_5 | CCCATATGTTTTCAGCCAAGTTCCATCTGTCGGCGAGTCC |
| adpARev_5 | GGACTCGCCGACAGATGGAACTTGGCTGAAAACATATGGGcctaccctacgtcctcctgc |
| adpAFor_6 | TCTGTCGGCGAGTCCTTGGCGGAACGGTTGCCCAACCCCG |
| adpARev_6 | CGGGGTTGGGCAACCGTTCCGCCAAGGACTCGCCGACAGAcctaccctacgtcctcctgc |
| adpAFor_7 | GGTTGCCCAACCCCGGCGCCGGCACCCGAGAATGCCCCGC |
| adpARev_7 | GCGGGGCATTCTCGGGTGCCGGCGCCGGGGTTGGGCAACCcctaccctacgtcctcctgc |
| adpAFor_8 | CCGAGAATGCCCCGCCACAAGGGCCCGCACCCCCCGCGTA |
| adpARev_8 | TACGCGGGGGGTGCGGGCCCTTGTGGCGGGGCATTCTCGGcctaccctacgtcctcctgc |
| adpAFor_9 | CGCACCCCCCGCGTACGGGGGCGTTCGCCACGCCGCCCCT |
| adpARev_9 | AGGGGCGGCGTGGCGAACGCCCCCGTACGCGGGGGGTGCGcctaccctacgtcctcctgc |
| adpAFor_10 | CGCCACGCCGCCCCTTCCCCGGGCGGCCATCGGACGGAAG |
| adpARev_10 | CTTCCGTCCGATGGCCGCCCGGGGAAGGGGCGGCGTGGCGcctaccctacgtcctcctgc |
| adpAFor_11 | GCCATCGGACGGAAGACTTCGCGATCGATCGCTTCACGCC |
| adpARev_11 | GGCGTGAAGCGATCGATCGCGAAGTCTTCCGTCCGATGGCcctaccctacgtcctcctgc |
| adpAFor_12 | CGATCGCTTCACGCCAAGTGGCCTTGTCGACAATCCACCG |
| adpARev_12 | CGGTGGATTGTCGACAAGGCCACTTGGCGTGAAGCGATCGcctaccctacgtcctcctgc |
| adpAFor_13 | GTCGACAATCCACCGGATGGAGAACTTGTCACGCCGGCGG |
| adpARev_13 | CCGCCGGCGTGACAAGTTCTCCATCCGGTGGATTGTCGACcctaccctacgtcctcctgc |
| adpAFor_14 | TTGTCACGCCGGCGGCACGGGACGCAGTAGATTCGATCAT |
| adpARev_14 | ATGATCGAATCTACTGCGTCCCGTGCCGCCGGCGTGACAAcctaccctacgtcctcctgc |
| adpAFor_15 | AGTAGATTCGATCATGGGTACCGAAGACTGGGGTCTCGTG |
| adpARev_15 | CACGAGACCCCAGTCTTCGGTACCCATGATCGAATCTACTcctaccctacgtcctcctgc |
| adpAFor_16 | GACTGGGGTCTCGTGCAAAACCGAGGGGAAACGTGCAGGA |
| adpARev_16 | TCCTGCACGTTTCCCCTCGGTTTTGCACGAGACCCCAGTCcctaccctacgtcctcctgc |
| adpAFor_17 | GGGAAACGTGCAGGAGCGACACGACCAGGGAGACGCGAAC |
| adpARev_17 | GTTCGCGTCTCCCTGGTCGTGTCGCTCCTGCACGTTTCCCcctaccctacgtcctcctgc |
| adpAFor_18 | CAGGGAGACGCGAACACCGAGGGGGGCTTAGCGTCATGAG |
| adpARev_18 | CTCATGACGCTAAGCCCCCCTCGGTGTTCGCGTCTCCCTGcctaccctacgtcctcctgc |
| adpAFor_19 | GCTTAGCGTCATGAGCCAGGACTCC |
| adpARev_19 | GGAGTCCTGGCTCATGACGCTAAGCcctaccctacgtcctcctgc |
| MtrAFor_1 | TCATATCGACATCATCCCATTTCCGTATCAGTCTCAAGGC |
| MtrARev_1 | GCCTTGAGACTGATACGGAAATGGGATGATGTCGATATGAcctaccctacgtcctcctgc |
| MtrAFor_2 | TATCAGTCTCAAGGCGGCTGGTGAGATACCTCACTGACCT |
| MtrARev_2 | AGGTCAGTGAGGTATCTCACCAGCCGCCTTGAGACTGATAcctaccctacgtcctcctgc |
| MtrAFor_3 | ATACCTCACTGACCTGCGGTGACGTCGGTGATCGCCCACC |
| MtrARev_3 | GGTGGGCGATCACCGACGTCACCGCAGGTCAGTGAGGTATcctaccctacgtcctcctgc |
| MtrAFor_4 | CGGTGATCGCCCACCGTCGACTGCCGGTGTCCGTGGGTGT |
| MtrARev_4 | ACACCCACGGACACCGGCAGTCGACGGTGGGCGATCACCGcctaccctacgtcctcctgc |
| MtrAFor_5 | GGTGTCCGTGGGTGTTGATGCCAGACATGGATGTCACCCC |
| MtrARev_5 | GGGGTGACATCCATGTCTGGCATCAACACCCACGGACACCcctaccctacgtcctcctgc |
| MtrAFor_6 | CATGGATGTCACCCCCGTGGGCCCCCGTGGTCGGCCTGCA |
| MtrARev_6 | TGCAGGCCGACCACGGGGGCCCACGGGGGTGACATCCATGcctaccctacgtcctcctgc |
| MtrAFor_7 | ACCCCCGTGGGCCCCCGTGGTCGGCCTGCACCGTACCCTG |
| MtrARev_7 | CAGGGTACGGTGCAGGCCGACCACGGGGGCCCACGGGGGTcctaccctacgtcctcctgc |
| dnaAFor_1 | TAGGTGGACGACGAGGAGCGGGCGACCGGCCCGGCGCCCT |
| dnaARev_1 | AGGGCGCCGGGCCGGTCGCCCGCTCCTCGTCGTCCACCTAcctaccctacgtcctcctgc |
| dnaAFor_2 | CCGGCCCGGCGCCCTCGGCGTACCGCGGTTGCGAAGTCCT |
| dnaARev_2 | AGGACTTCGCAACCGCGGTACGCCGAGGGCGCCGGGCCGGcctaccctacgtcctcctgc |
| dnaAFor_3 | CGGTTGCGAAGTCCTCGCGCCGCCTCAGCCGATTGTCGGT |
| dnaARev_3 | ACCGACAATCGGCTGAGGCGGCGCGAGGACTTCGCAACCGcctaccctacgtcctcctgc |
| dnaAFor_4 | CAGCCGATTGTCGGTAGGCAGCACGTCATGACCCGTTTAG |
| dnaARev_4 | CTAAACGGGTCATGACGTGCTGCCTACCGACAATCGGCTGcctaccctacgtcctcctgc |
| dnaAFor_5 | TCATGACCCGTTTAGCGGATCAGGCGGACAGCTCAGAGCG |
| dnaARev_5 | CGCTCTGAGCTGTCCGCCTGATCCGCTAAACGGGTCATGAcctaccctacgtcctcctgc |
| dnaAFor_6 | GGACAGCTCAGAGCGACCCTTGGAACGACGGTTCGCAAGG |
| dnaARev_6 | CCTTGCGAACCGTCGTTCCAAGGGTCGCTCTGAGCTGTCCcctaccctacgtcctcctgc |
| dnaAFor_7 | CGACGGTTCGCAAGGATGGCTCGGCCGGCACGCGTACGCA |
| dnaARev_7 | TGCGTACGCGTGCCGGCCGAGCCATCCTTGCGAACCGTCGcctaccctacgtcctcctgc |
| dnaAFor_8 | CGGCACGCGTACGCATCCGCAGGCGGAAGCCGTGGGTCTT |
| dnaARev_8 | AAGACCCACGGCTTCCGCCTGCGGATGCGTACGCGTGCCGcctaccctacgtcctcctgc |
| dnaAFor_9 | GAAGCCGTGGGTCTTGGCGCGACGACGGTTGTTCGGCTGG |
| dnaARev_9 | CCAGCCGAACAACCGTCGTCGCGCCAAGACCCACGGCTTCcctaccctacgtcctcctgc |
| dnaAFor_10 | CGGTTGTTCGGCTGGAAGGTGCGCTTGCTCACTCGGGGGC |
| dnaARev_10 | GCCCCCGAGTGAGCAAGCGCACCTTCCAGCCGAACAACCGcctaccctacgtcctcctgc |
| dnaAFor_11 | TGCTCACTCGGGGGCTCCAGAAATGATTCGTAGATGGCGG |
| dnaARev_11 | CCGCCATCTACGAATCATTTCTGGAGCCCCCGAGTGAGCAcctaccctacgtcctcctgc |
| dnaAFor_12 | ATTCGTAGATGGCGGGACATCGCCTGGCTGTCACCGTGCG |
| dnaARev_12 | CGCACGGTGACAGCCAGGCGATGTCCCGCCATCTACGAATcctaccctacgtcctcctgc |
| dnaAFor_13 | GGCTGTCACCGTGCGCCCACGAGTAGCTCGCAATACGCCC |
| dnaARev_13 | GGGCGTATTGCGAGCTACTCGTGGGCGCACGGTGACAGCCcctaccctacgtcctcctgc |
| dnaAFor_14 | GCTCGCAATACGCCCGAGTGCACCGCTTCACGATCACTGA |
| dnaARev_14 | TCAGTGATCGTGAAGCGGTGCACTCGGGCGTATTGCGAGCcctaccctacgtcctcctgc |
| dnaAFor_15 | CTTCACGATCACTGACCGTGATCTTTGCCCATCGGAGGCA |
| dnaARev_15 | TGCCTCCGATGGGCAAAGATCACGGTCAGTGATCGTGAAGcctaccctacgtcctcctgc |
| dnaAFor_16 | TGCCCATCGGAGGCAGGCGGCAGCAGCCATCGACAACTCG |
| dnaARev_16 | CGAGTTGTCGATGGCTGCTGCCGCCTGCCTCCGATGGGCAcctaccctacgtcctcctgc |
| dnaAFor_17 | GCCATCGACAACTCGACCTGGTTACGGTACGCGCGGCTAC |
| dnaARev_17 | GTAGCCGCGCGTACCGTAACCAGGTCGAGTTGTCGATGGCcctaccctacgtcctcctgc |
| dnaAFor_18 | GGTACGCGCGGCTACGCCATCCGGTCAAACCGACCTGTCG |
| dnaARev_18 | CGACAGGTCGGTTTGACCGGATGGCGTAGCCGCGCGTACCcctaccctacgtcctcctgc |
| dnaAFor_19 | CAAACCGACCTGTCGCCACCCCCCATTGTGCACAGGCTGT |
| dnaARev_19 | ACAGCCTGTGCACAATGGGGGGTGGCGACAGGTCGGTTTGcctaccctacgtcctcctgc |
| dnaAFor_20 | TTGTGCACAGGCTGTGGACAACAACTTGAACCACGTCATC |
| dnaARev_20 | GATGACGTGGTTCAAGTTGTTGTCCACAGCCTGTGCACAAcctaccctacgtcctcctgc |
| dnaAFor_21 | TTGAACCACGTCATCCGGCGCGACTACCGTGGATGGACTC |
| dnaARev_21 | GAGTCCATCCACGGTAGTCGCGCCGGATGACGTGGTTCAAcctaccctacgtcctcctgc |
| dnaAFor_22 | ACCGTGGATGGACTCCACAATCTTTTCCGTTCTGTCCTTA |
| dnaARev_22 | TAAGGACAGAACGGAAAAGATTGTGGAGTCCATCCACGGTcctaccctacgtcctcctgc |
| dnaAFor_23 | TCCGTTCTGTCCTTACCTGTCCTCACGGGTTCTGTCCTCA |
| dnaARev_23 | TGAGGACAGAACCCGTGAGGACAGGTAAGGACAGAACGGAcctaccctacgtcctcctgc |
| dnaAFor_24 | CGGGTTCTGTCCTCACGGACATCGACCCACCGTCCCCGAG |
| dnaARev_24 | CTCGGGGACGGTGGGTCGATGTCCGTGAGGACAGAACCCGcctaccctacgtcctcctgc |
| dnaAFor_25 | CCCACCGTCCCCGAGAACCACACCATCAGGGGACCTGCGA |
| dnaARev_25 | TCGCAGGTCCCCTGATGGTGTGGTTCTCGGGGACGGTGGGcctaccctacgtcctcctgc |
| dnaAFor_26 | TCAGGGGACCTGCGAGAAAGCGTGCCCTGTGGCTGACGTT |
| dnaARev_26 | AACGTCAGCCACAGGGCACGCTTTCTCGCAGGTCCCCTGAcctaccctacgtcctcctgc |
| dnaAFor_27 | CCTGTGGCTGACGTTCCTGCTGATCTTGCCGCAG |
| dnaARev_27 | CTGCGGCAAGATCAGCAGGAACGTCAGCCACAGGcctaccctacgtcctcctgc |
| bldMFor_1 | TTGCGCTTGTCCCCTTGAAACCGTGTCGAACGGATAGGTG |
| bldMRev_1 | CACCTATCCGTTCGACACGGTTTCAAGGGGACAAGCGCAAcctaccctacgtcctcctgc |
| bldMFor_2 | TCGAACGGATAGGTGGGCGGGTGGATGGCTGATGGCCGGG |
| bldMRev_2 | CCCGGCCATCAGCCATCCACCCGCCCACCTATCCGTTCGAcctaccctacgtcctcctgc |
| bldMFor_3 | TGGCTGATGGCCGGGCTCCTCGGGGCCCCGCCGGTCCGGG |
| bldMRev_3 | CCCGGACCGGCGGGGCCCCGAGGAGCCCGGCCATCAGCCAcctaccctacgtcctcctgc |
| bldMFor_4 | CCCCGCCGGTCCGGGTCGGTACACCTACTGTCTAAGTAGA |
| bldMRev_4 | TCTACTTAGACAGTAGGTGTACCGACCCGGACCGGCGGGGcctaccctacgtcctcctgc |
| bldMFor_5 | TACTGTCTAAGTAGATGTAAATATGACTCATTGCGAATCT |
| bldMRev_5 | AGATTCGCAATGAGTCATATTTACATCTACTTAGACAGTAcctaccctacgtcctcctgc |
| bldMFor_6 | ACTCATTGCGAATCTAGCCACAGACACCGCGAAAAGGGAA |
| bldMRev_6 | TTCCCTTTTCGCGGTGTCTGTGGCTAGATTCGCAATGAGTcctaccctacgtcctcctgc |
| bldMFor_7 | ACCGCGAAAAGGGAAGAAAACCCGCTAAATGGGGCATAGC |
| bldMRev_7 | GCTATGCCCCATTTAGCGGGTTTTCTTCCCTTTTCGCGGTcctaccctacgtcctcctgc |
| bldMFor_8 | TAAATGGGGCATAGCTTTCGATGAACGACAGAAGGCTCGC |
| bldMRev_8 | GCGAGCCTTCTGTCGTTCATCGAAAGCTATGCCCCATTTAcctaccctacgtcctcctgc |
| bldMFor_9 | CGACAGAAGGCTCGCGTCGCTCTCCTCTGTACGCGCCCCC |
| bldMRev_9 | GGGGGCGCGTACAGAGGAGAGCGACGCGAGCCTTCTGTCGcctaccctacgtcctcctgc |
| bldMFor_10 | TCTGTACGCGCCCCCTCACGTAGAGTGCCGAAGGGTGCCG |
| bldMRev_10 | CGGCACCCTTCGGCACTCTACGTGAGGGGGCGCGTACAGAcctaccctacgtcctcctgc |
| bldMFor_11 | TGCCGAAGGGTGCCGTCCGACCCGTAACTCTTTCGAGTGA |
| bldMRev_11 | TCACTCGAAAGAGTTACGGGTCGGACGGCACCCTTCGGCAcctaccctacgtcctcctgc |
| bldMFor_12 | AACTCTTTCGAGTGACCGTCGTTGAGAGTGCGGAGGCGGT |
| bldMRev_12 | ACCGCCTCCGCACTCTCAACGACGGTCACTCGAAAGAGTTcctaccctacgtcctcctgc |
| bldMFor_13 | GAGTGCGGAGGCGGTTGAAGGAACAAGCGATCGGGCAGGT |
| bldMRev_13 | ACCTGCCCGATCGCTTGTTCCTTCAACCGCCTCCGCACTCcctaccctacgtcctcctgc |
| bldMFor_14 | AGCGATCGGGCAGGTGTCCGAGAGCGTCAATCGCACAGGT |
| bldMRev_14 | ACCTGTGCGATTGACGCTCTCGGACACCTGCCCGATCGCTcctaccctacgtcctcctgc |
| bldMFor_15 | GTCAATCGCACAGGTGACGATTACGTACAGCCCTGGAGGC |
| bldMRev_15 | GCCTCCAGGGCTGTACGTAATCGTCACCTGTGCGATTGACcctaccctacgtcctcctgc |
| bldMFor_16 | TACAGCCCTGGAGGCTCAAGGTGACGCGCTACAGCTGCGA |
| bldMRev_16 | TCGCAGCTGTAGCGCGTCACCTTGAGCCTCCAGGGCTGTAcctaccctacgtcctcctgc |
| bldMFor_17 | GCGCTACAGCTGCGAGAGCCGCGGAGGTCAGGCATGACAT |
| bldMRev_17 | ATGTCATGCCTGACCTCCGCGGCTCTCGCAGCTGTAGCGCcctaccctacgtcctcctgc |
| bldMFor_18 | GGTCAGGCATGACATCCGTTCTCG |
| bldMRev_18 | CGAGAACGGATGTCATGCCTGACCcctaccctacgtcctcctgc |
| filPFor_1 | GGGTAGCCACTCAAAAGGCTGGACATTCTCCAGATCAAAC |
| filPRev_1 | GTTTGATCTGGAGAATGTCCAGCCTTTTGAGTGGCTACCCcctaccctacgtcctcctgc |
| filPFor_2 | TTCTCCAGATCAAACGGGCATACGCTCGATGACACGCCGC |
| filPRev_2 | GCGGCGTGTCATCGAGCGTATGCCCGTTTGATCTGGAGAAcctaccctacgtcctcctgc |
| filPFor_3 | TCGATGACACGCCGCTTCGGCCCCTAGGATTCCCTCTAAC |
| filPRev_3 | GTTAGAGGGAATCCTAGGGGCCGAAGCGGCGTGTCATCGAcctaccctacgtcctcctgc |
| filPFor_4 | AGGATTCCCTCTAACACCTCACCGGTCTCATTCGACAGGA |
| filPRev_4 | TCCTGTCGAATGAGACCGGTGAGGTGTTAGAGGGAATCCTcctaccctacgtcctcctgc |
| filPFor_5 | TCTCATTCGACAGGAAACCCATGAG |
| filPRev_5 | CTCATGGGTTTCCTGTCGAATGAGAcctaccctacgtcctcctgc |
| whiIFor_1 | AGGGGCTGTTTTCGGCCTGTTCCGGTGTCCCTCGTGGGGG |
| whiIRev_1 | CCCCCACGAGGGACACCGGAACAGGCCGAAAACAGCCCCTcctaccctacgtcctcctgc |
| whiIFor_2 | TGTCCCTCGTGGGGGTGACGGAGCGGGGGCCGCTCGTAGG |
| whiIRev_2 | CCTACGAGCGGCCCCCGCTCCGTCACCCCCACGAGGGACAcctaccctacgtcctcctgc |
| whiIFor_3 | GGGGCCGCTCGTAGGATGATCGATCACGCGTCCGAATTGC |
| whiIRev_3 | GCAATTCGGACGCGTGATCGATCATCCTACGAGCGGCCCCcctaccctacgtcctcctgc |
| whiIFor_4 | ACGCGTCCGAATTGCCCTAATTGTTACTCACCAAATCGTG |
| whiIRev_4 | CACGATTTGGTGAGTAACAATTAGGGCAATTCGGACGCGTcctaccctacgtcctcctgc |
| whiIFor_5 | ACTCACCAAATCGTGATCTTTCCCTAAAGGCGGGCGGCCC |
| whiIRev_5 | GGGCCGCCCGCCTTTAGGGAAAGATCACGATTTGGTGAGTcctaccctacgtcctcctgc |
| whiIFor_6 | AAAGGCGGGCGGCCCAGCTGCCGAAGGAGTCAGTGACCCC |
| whiIRev_6 | GGGGTCACTGACTCCTTCGGCAGCTGGGCCGCCCGCCTTTcctaccctacgtcctcctgc |
| whiIFor_7 | GGAGTCAGTGACCCCTTCACAGCACGGGTTCGCCCCCGGC |
| whiIRev_7 | GCCGGGGGCGAACCCGTGCTGTGAAGGGGTCACTGACTCCcctaccctacgtcctcctgc |
| whiIFor_8 | GGGTTCGCCCCCGGCTTCTTCCCCGAGCCGGCTCCGTCCC |
| whiIRev_8 | GGGACGGAGCCGGCTCGGGGAAGAAGCCGGGGGCGAACCCcctaccctacgtcctcctgc |
| whiIFor_9 | AGCCGGCTCCGTCCCGCACCCTTCCCCCCAGGAGGCCTGG |
| whiIRev_9 | CCAGGCCTCCTGGGGGGAAGGGTGCGGGACGGAGCCGGCTcctaccctacgtcctcctgc |
| whiIFor_10 | CCCCAGGAGGCCTGGTGTCCGTTCT |
| whiIRev_10 | AGAACGGACACCAGGCCTCCTGGGGcctaccctacgtcctcctgc |
| Binding_site_T1A | CCCTCACCCCCCACCCCTCACTCGTGTTCCCCGCGCGCTC |
| Binding_site_T1A | GAGCGCGCGGGGAACACGAGTGAGGGGTGGGGGGTGAGGGcctaccctacgtcctcctgc |
| Binding_site_C4G | CCCTCACCCCCCACCCCTCACTCCTGATCCCCGCGCGCTC |
| Binding_site_C4G | GAGCGCGCGGGGATCAGGAGTGAGGGGTGGGGGGTGAGGGcctaccctacgtcctcctgc |
| Binding_site_BOTH | CCCTCACCCCCCACCCCTCACTCCTGTTCCCCGCGCGCTC |
| Binding_site_BOTH | GAGCGCGCGGGGAACAGGAGTGAGGGGTGGGGGGTGAGGGcctaccctacgtcctcctgc |

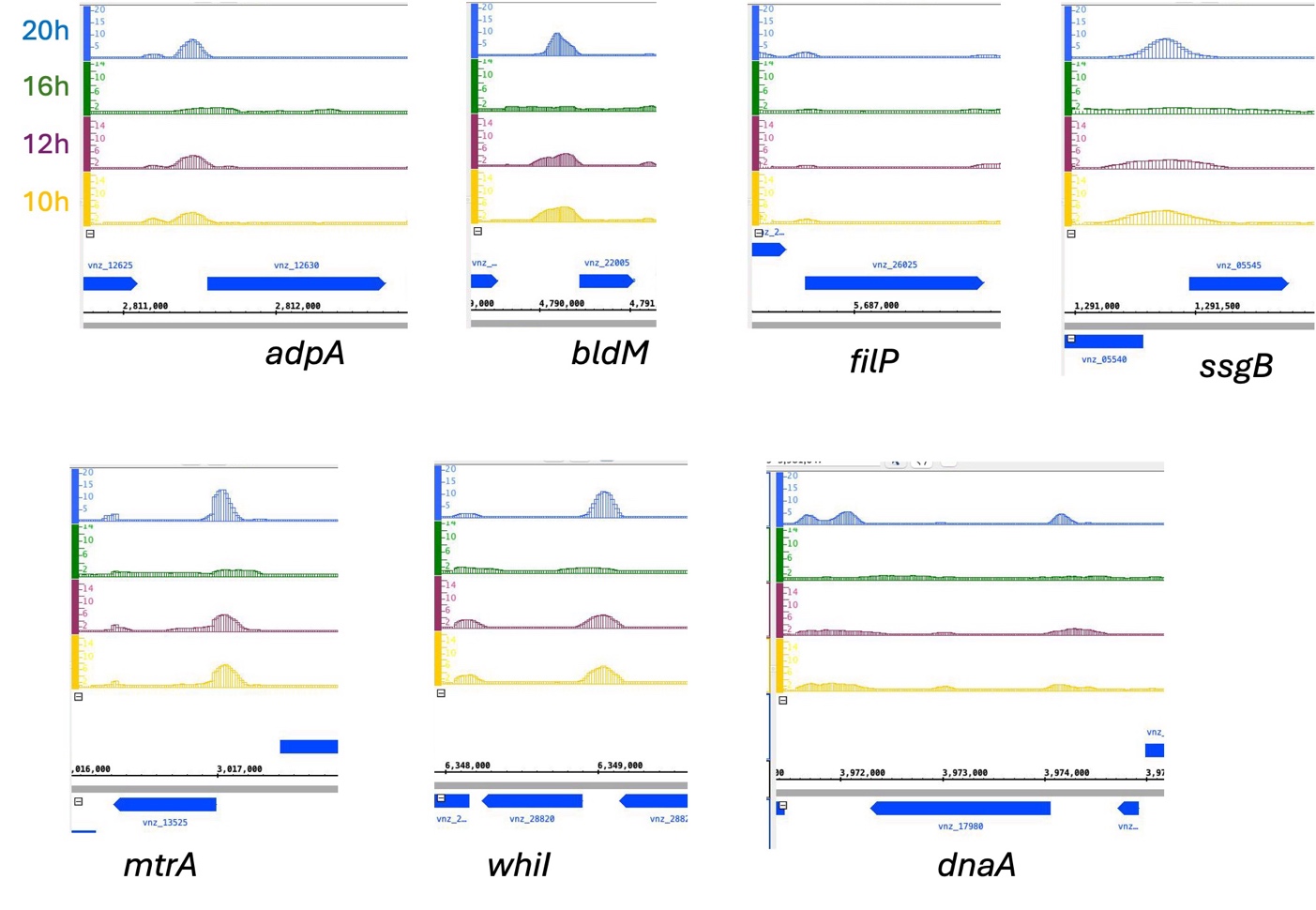

**Figure S1. ChIP-Seq peaks for key developmental genes in *S. venezualae*.** MtrA-FLAG binds promoters of key developmental genes in *S. venezualae* (Som *et al.*, 2017)
